# MorphoNavigator-3D: Generalizable single-cell phenotyping of cancer spheroids using Bayesian-optimized deep-learning workflows

**DOI:** 10.1101/2024.09.08.611898

**Authors:** Isabel Mogollon, Michaela Feodoroff, Anna Nylund, Alba Montedeoca, Gantugs Atarsaikhan, Pedro Neto, Peter Horvath, Antti Rannikko, Vincenzo Cerullo, Vilja Pietiäinen, Lassi Paavolainen

**Author notes:** These authors equally share second authorship (Michaela Feodoroff, Anna Nylund). These authors jointly supervised this work (Vilja Pietiäinen, Lassi Paavolainen). Correspondence: Isabel Mogollon, and Lassi Paavolainen, Tukholmankatu 8, 00290, Helsinki, Finland.

## Abstract

Accurate quantification of drug responses in 3D tumor–immune co-cultures remains challenging because complex spatial architecture and cellular heterogeneity limit the interpretability of bulk viability assays. Here, we present **MorphoNavigator-3D** (“**Morpho**logical **Navigator** in **3D**”;**MoNa-3D**), an automated framework for high-resolution, annotation-free single-cell analysis in complex 3D co-cultures. The approach integrates optimized live-cell staining, deep learning–based segmentation, and Bayesian optimization (BO) to adapt end-to-end image-analysis workflows across diverse experimental conditions.

MoNa-3D was applied to clear cell renal cell carcinoma (ccRCC)–immune cell 3D-spheroid co-cultures, exposed to PI3K/mTOR pathway inhibitors and immunomodulatory compounds in a high-content imaging-based drug screen. The pipeline was used to extract multiscale phenotypic features encompassing ATP-based cell viability, morphology, nuclear remodeling, spatial dispersion, and immune infiltration. This analysis resolved distinct drug-induced phenotypes: PI3K/mTOR inhibitors promoted spheroid disintegration, nuclear enlargement, and immune exclusion, whereas immunomodulators preserved spheroid architecture and T-cell engagement. Multivariate phenotypic integration distinguished drug classes and revealed intra-class variation, including divergent spatial responses to dual PI3K/mTOR versus mTORC1 inhibition. These phenotypes were consistent with known drug mechanisms, supporting the biological interpretability of the framework. Together, these findings establish MoNa-3D as a generalizable platform for multidimensional phenotypic profiling across complex 3D multicellular systems, supporting applications in drug discovery, tumor–immune interaction studies, and precision oncology.

**JOURNAL HIGHLIGHTS:**

- Development of MorphoNavigator-3D (MoNa-3D), a fully automated 3D image analysis framework combining deep learning segmentation with Bayesian optimization.
  - Enables multidimensional single-cell phenotyping in tumor–immune spheroid co-cultures under diverse drug conditions.
  - Resolves structural disintegration, nuclear remodeling, and spatial immune engagement across drug classes.
  - Detects subtle intra-class differences between PI3K/mTOR pathway inhibitors.
  - Provides a scalable platform for drug discovery, immunotherapy research, and precision oncology.

## INTRODUCTION

Three-dimensional (3D) cell cultures have become essential for studying cancer biology in physiologically relevant models^1^. Unlike 2D cultures, 3D cultures, such as spheroids and organoids, better mimic tumor heterogeneity, microenvironmental complexity, and drug responses^2,3^. Co-cultures of tumor and immune cells are increasingly used to model tumor–immune interactions *in vitro*, including those between cancer cells and CD8⁺/CD4⁺ T-cells, regulatory T-cells, NK cells, and myeloid-derived suppressor cells^4,5^. Simultaneously, the limited clinical translation of promising therapies highlights a critical need for improved preclinical models and single-cell level analytics that accurately recapitulate tumor–immune interactions and drug responses. However, single-cell analysis in such systems remains technically challenging, especially for high-throughput (HT) experiments, although recent light-sheet datasets have demonstrated the feasibility of high-resolution imaging in multicultures^6^.

Among solid tumors, clear cell renal cell carcinoma (ccRCC) represents a particularly challenging model due to its extensive intratumoral heterogeneity and strong microenvironmental influence. Furthermore, ccRCC is highly immunogenic and frequently exhibits substantial immune infiltration, although immune cell function is often constrained by metabolic and immunosuppressive features of the tumor microenvironment^7,8^,.

Segmenting individual cells in spheroids is challenging due to dense clustering of cells and anisotropic image resolution and depth-dependent light scattering. Manual annotation is impractical at scale, while conventional deep-learning (DL)^9^ approaches require large annotated datasets and struggle to generalize across diverse staining or imaging conditions^10–12^. Semi-automatic tools have improved efficiency in generating 2D and 3D labels^13^, but scalability remains a bottleneck in large phenotypic screens. While Bayesian optimization offers a powerful strategy to tune segmentation workflows in a data-driven, annotation-free manner^14^, it remains underused in this context.

Beyond segmentation, robust phenotyping in 3D models requires careful dye selection to ensure deep tissue penetration, low toxicity, and cell-type-specific labelling^15–17^. This is critical for distinguishing different cell types, such as immune and cancer cell populations in co-culture settings. Moreover, traditional viability assays alone fail to capture morphological and spatial drug effects, such as nuclear remodeling, spheroid disintegration, or immune exclusion, that are often important for interpreting therapeutic responses^17,18^. Spatial metrics, including immune infiltration or exclusion, are emerging as key indicators of drug response, particularly in the context of immunomodulation^19–21^. However, only few tools currently integrate such spatial features into single-cell 3D analysis pipelines^6,22^.

To address these gaps, we developed MorphoNavigator-3D (MoNa-3D), an integrated framework that combines optimized staining protocols, DL-based segmentation, and Bayesian optimization for automated, high-resolution single-cell analysis in 3D co-cultures. This framework provides a scalable approach for multidimensional phenotyping of co-cultures, enabling the investigation of how perturbations influence cellular traits by considering cell viability, morphology, and spatial arrangement. In our proof-of-concept study, we applied MoNa-3D to ccRCC–T-cell co-cultures, modeling aspects of the immune-rich ccRCC tumor microenvironment^7,8^, and further evaluates its applicability in a patient-derived ccRCC–PBMC model. Our results demonstrate that MoNa-3D resolves both inter-and intra-class drug responses, revealing distinct effects on spheroid architecture, immune engagement, and spatial organization. In particular, PI3K/mTOR inhibitors induced structural disintegration, nuclear enlargement and immune exclusion, whereas immunomodulators preserved spheroid integrity and T-cell engagement. Together, these findings demonstrate that MoNa-3D, by enabling multidimensional single-cell phenotyping in complex 3D tumor–immune systems, can reveal biologically meaningful treatment responses that are not captured by conventional viability measurements.

## RESULTS

### Optimized fluorescent live dyes enable visualization of individual cells in co-culture 3D spheroids

The MoNa-3D workflow integrates optimized cell labelling and high-content imaging with DL-based segmentation, Bayesian optimization, and downstream phenotypic profiling (Fig. 1). Spheroids were cultured for 3 days, consistent with established Drug Sensitivity and Resistance Testing (DSRT) conditions^23–27^, before 3D imaging and quantitative single-cell analysis.

**Figure 1.**
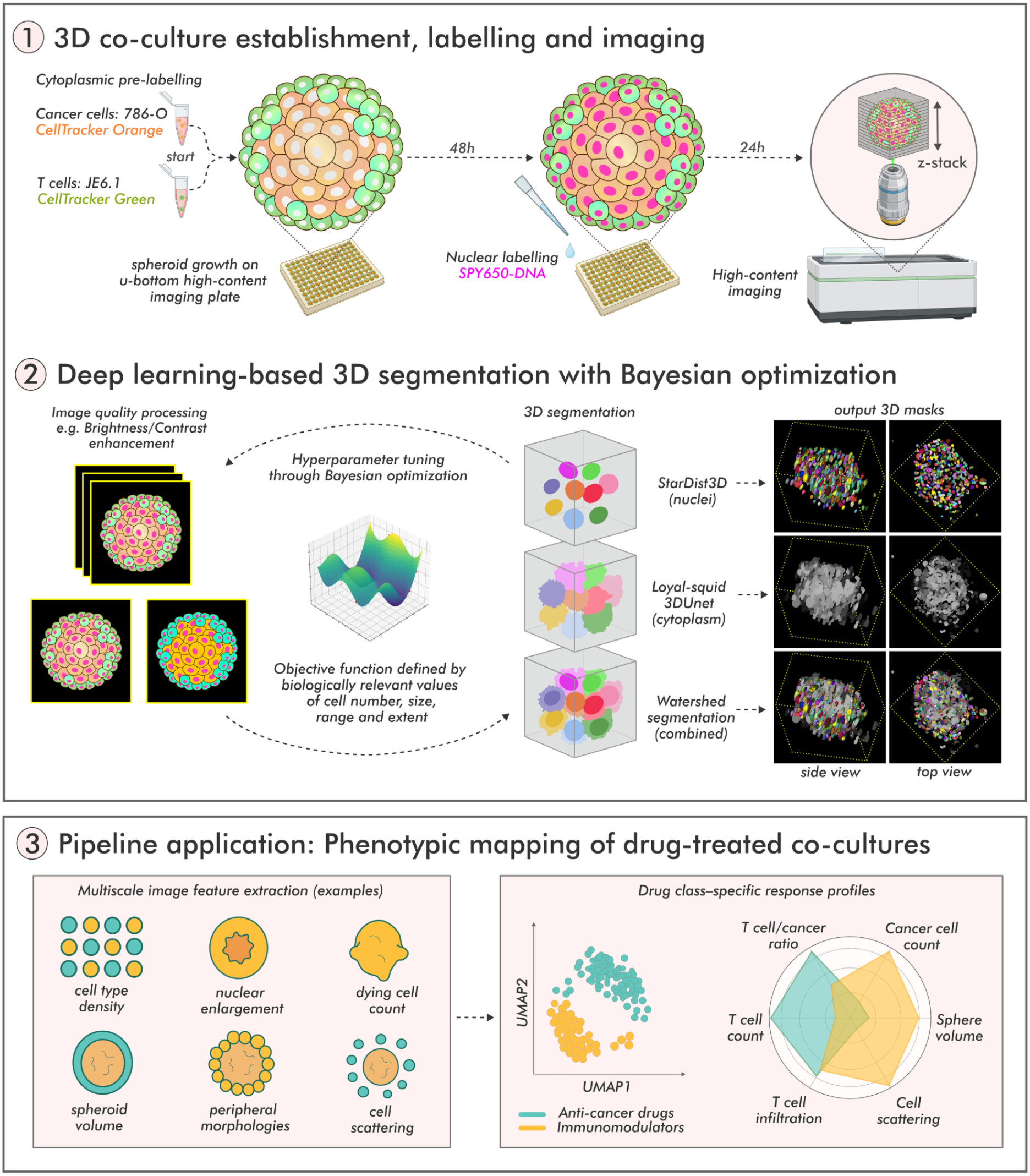
Overview of the integrated wet-lab and dry-lab workflow for segmentation optimization in 3D cancer–immune co-cultures in MorphoNavigator-3D, followed by application to drug-treated spheroids. (1) Renal cancer cell (786-O) and T-cell (JE6.1) populations were pre-labelled with CellTracker Orange and CellTracker Green, respectively, and seeded into 3D co-cultures in 384-well plates (500:250 ratio). After 48 h, spheroids were stained with SPY650-DNA to enable nuclear visualization for segmentation. At 72 h, 3D stack images of whole spheroids were acquired with a high-content fluorescence confocal microscope. (2) Acquired image stacks underwent preprocessing, which included multiple steps, e.g., channel merging and brightness/contrast enhancement. Nuclear and cell segmentation was performed using multiple DL-based approaches (StarDist3D, ‘loyal-squid’ 3D U-Net), and watershed-based methods. A Bayesian optimization framework was used to refine preprocessing and segmentation parameters for improved accuracy, guided by biologically meaningful metrics (e.g., cell count, cell size, and spatial extent). Output 3D masks shown in the figure were rendered in Napari. (3) Schematic overview of a Python-based pipeline for single-cell and multicellular feature extraction from drug-treated spheroids and downstream analysis revealing drug class–specific response patterns.

To achieve accurate single-cell segmentation, the imaging conditions were optimized for non-toxic combinations of live dyes, concentrations, and incubation times. The primary model consisted of 3D spheroids of the ccRCC cell line 786-O, which recapitulates many genetic and phenotypic characteristics of ccRCC and is widely used as a preclinical model^28^. First, we sought to identify a far-red nuclear dye that maximized light penetration and fluorescence intensity within 3D spheroids. Three live-cell nuclear dyes—NucSpot, SYTO, and SPY650—were evaluated (Methods), with SPY650 exhibiting approximately 50% and 300% higher fluorescence intensity than SYTO and NucSpot, respectively, together with deeper penetration (Fig. 2A, Supp. Fig. 1A). SPY650 incubation was then optimized by testing three conditions: 72 h, 24 h, and 1 h before imaging. Compared with the 24 h incubation, the 1 h incubation achieved only 27.3–47.3% of the fluorescence intensity, depending on the culture model (Fig. 2A), whereas 72 h reduced fluorescence to 18.0% of the 24 h signal and caused partial spheroid disintegration, indicating dye-associated toxicity (Supp. Fig. 1A). CTG measurements confirmed that a 24 h SPY650 incubation did not reduce ATP-based cell viability compared with untreated controls or the 1 h condition (Fig. 2B).

**Figure 2.**
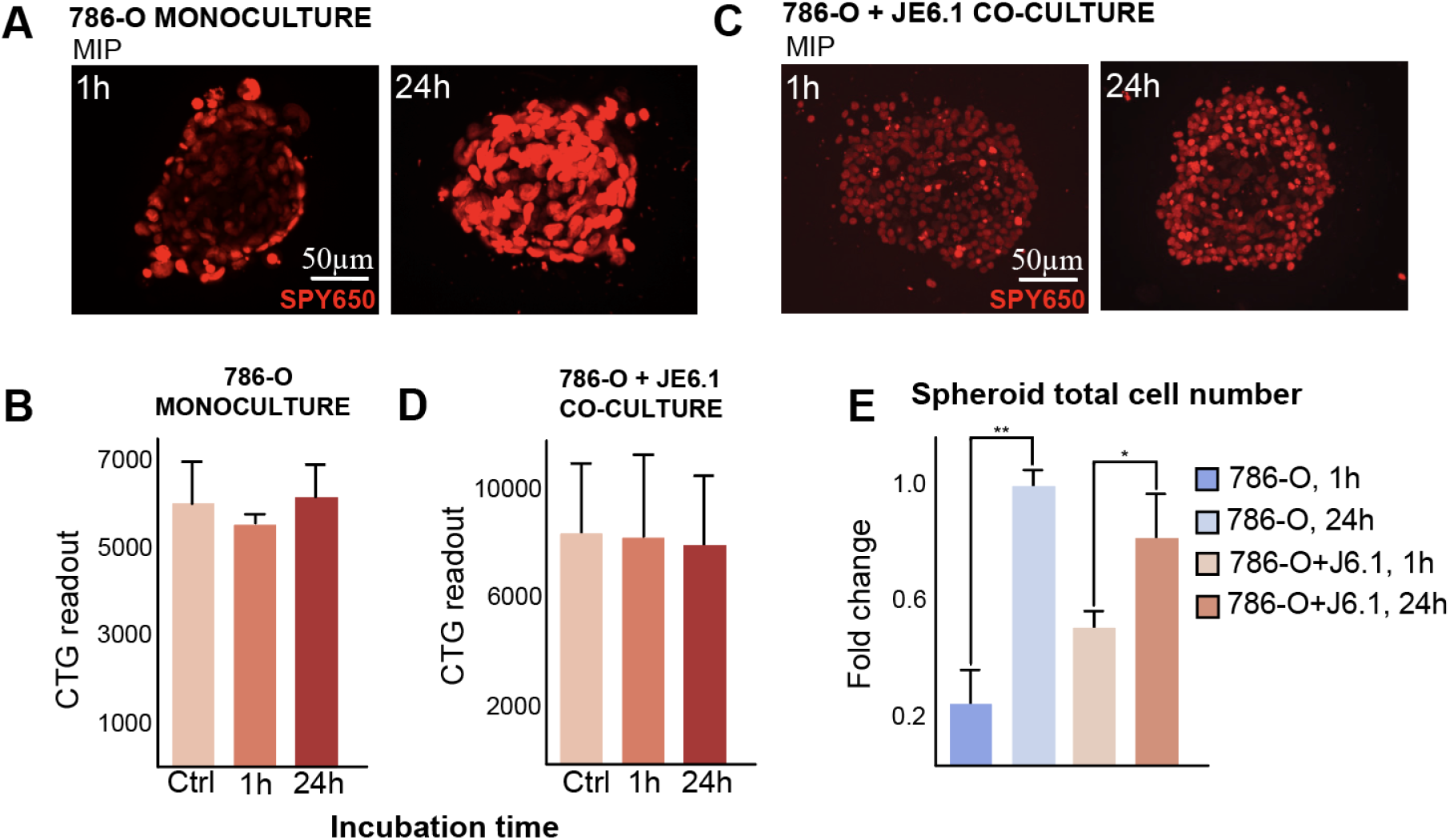
Optimization of SPY650 nuclear dye for 3D spheroid imaging and segmentation. (A) Representative maximum intensity projections (MIPs) showing nuclear staining of 786-O renal cancer spheroids with SPY650 after 1 h and 24 h incubation. The 24 h condition provided a stronger and more homogeneous nuclear signal. (B) Quantification of metabolic activity using CellTiter-Glo (CTG) assay in 786-O spheroids under control, 1 h, and 24 h SPY650 staining conditions, showing no reduction in ATP-based viability. (C) Application of 1 h and 24 h SPY650 protocols in 3D co-cultures of 786-O and JE6.1 cells. Clear nuclear signals are observed in both conditions, with improved clarity after 24 h incubation. (D) CTG viability measurements in co-cultures reveal no toxicity associated with SPY650 staining at either incubation time. (E) Quantification of segmented nuclei across conditions reveals a significant increase in detected cells under the 24 h SPY650 condition in both mono-and co-cultures, supporting improved nuclear segmentation performance. Data are normalized to the 786-O, 24 h condition (value = 1.0). *Scale bars: 50 µm.* Error bars indicate mean ± SEM; *p < 0.05, **p < 0.01.

The optimized 24 h SPY650 protocol was subsequently evaluated in 786-O/JE6.1 co-cultures, producing clear nuclear labeling of both cell types without affecting ATP-based viability (Fig. 2C,D). This condition also improved 3D single-cell nuclear segmentation, increasing the number of successfully segmented cells relative to the 1 h incubation (Fig. 2E). These findings were further supported by LIVE/DEAD assay cell staining, which indicated minimal cell death (Supp. Fig. 1B). Finally, SPY650 was compared with Hoechst as an alternative nuclear dye. Whereas SPY650 did not affect viability, Hoechst reduced CTG metabolic activity in co-cultures by approximately 98–99% relative to controls (Supp. Fig. 1C), indicating incompatibility with this spheroid system. Together, these results established SPY650 as the nuclear dye for all subsequent 3D segmentation experiments due to its higher fluorescence intensity, deeper penetration, and compatibility with live-cell imaging.

Next, we optimized cytoplasmic staining to achieve clear visualization of whole cells. We tested PKH26, CellVue, and CellTracker dyes in monocultures and co-cultures of 786-O cells with immune and endothelial cells (Methods). An important aspect of our protocol is the pre-labelling of cells before seeding them in U-bottom wells, ensuring uniform dye uptake throughout the 3D structure. Among the tested dyes, CellTracker produced the most consistent and homogeneous cytoplasmic staining (Fig. 3), whereas the lipophilic membrane dyes PKH26 and CellVue exhibited a more uneven, punctate labeling pattern under the tested conditions (Supp. Fig. 1D). Cells were then pre-labelled for 45 min with CellTracker at either 1 µM (1X) or 2 µM (2X) before spheroid formation to evaluate signal quality after 3 days of culture. The 2 µM (2X) concentration produced a stronger fluorescence signal without affecting ATP-based cell viability (Fig. 3A–C). In co-cultures, spectrally distinct CellTracker dyes enabled clear discrimination between cancer and immune cells (Fig. 3D). Combining SPY650 for nuclear staining with CellTracker dyes for cytoplasmic labeling did not alter CTG readouts compared with the corresponding CellTracker-only conditions and enabled comprehensive visualization of both cell types (Fig. 3D,E; Supp. Fig. 1E). Reducing the CellTracker Green concentration in JE6.1 cells to 0.5X further increased CTG readouts and improved T-cell visibility (Fig. 3E,F), suggesting a balance between dye concentration, viability, and image quality. These findings were supported by DRAQ7 staining, which detected only occasional DRAQ7-positive cells (Supp. Fig. 1E).

**Figure 3.**
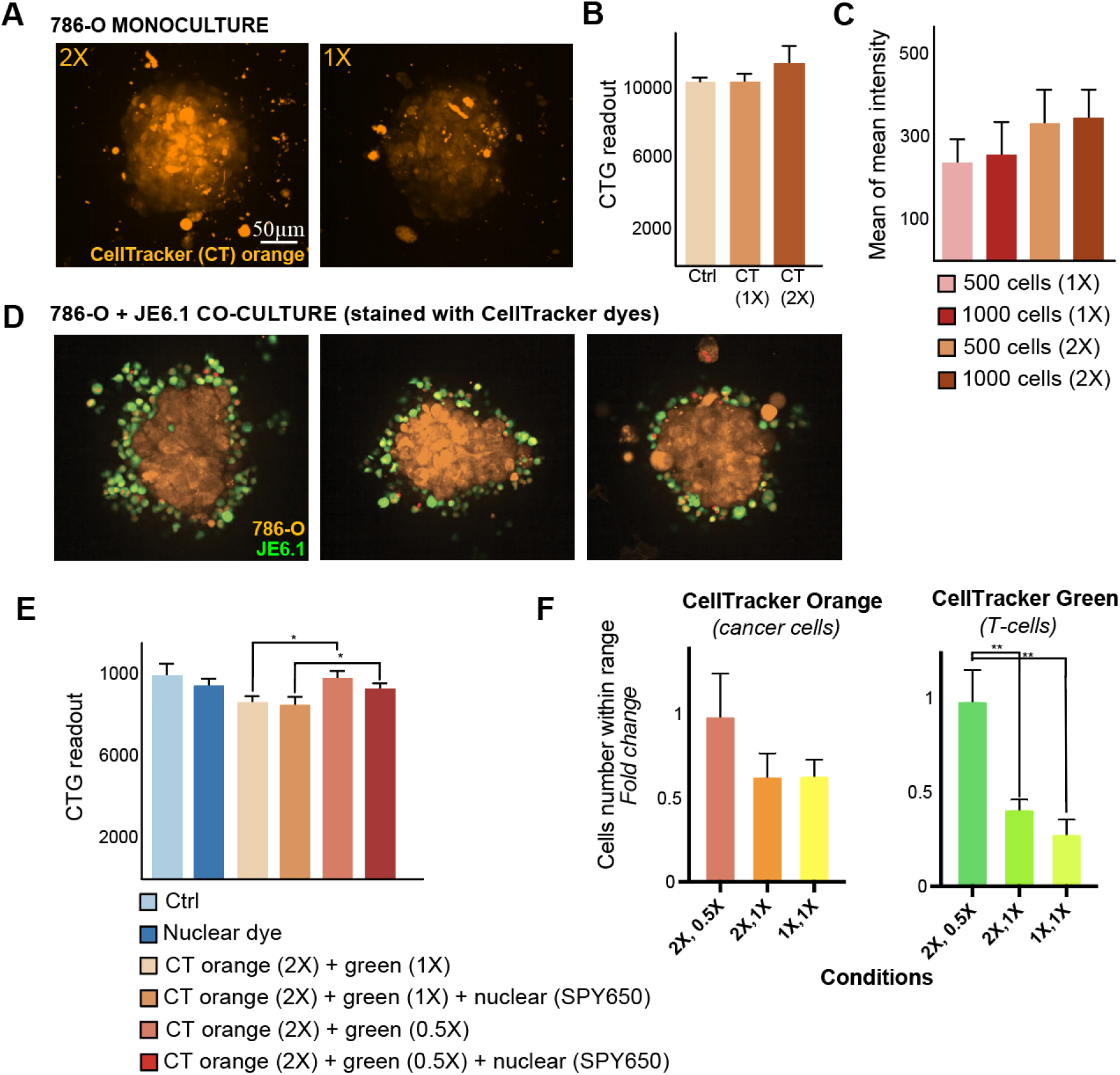
Optimization of CellTracker cytoplasmic dyes for clear cell visualization in 3D spheroids and co-cultures. (A) Representative MIPs of 786-O spheroids pre-labelled with CellTracker Orange (1X = 1 µM, 2X = 2 µM for 45 min) on day 0, followed by 72 h of spheroid growth at 37 C. The 2X concentration resulted in a more intense and uniform cytoplasmic signal. (B) ATP-based cell viability assessed by CellTiter-Glo (CTG) assay showed no cytotoxic effect at either concentration of the dye. (C) Quantification of mean CellTracker fluorescence intensity across conditions demonstrated higher signal in spheroids (500 or 1000 cells seeded and stained with 2X CellTracker) confirming dose-and density-dependent enhancement of cytoplasmic labelling. (D) Application of CellTracker Orange (2X) for 786-O cancer cells and CellTracker Green at varying concentrations for JE6.1 T-cells enabled reliable identification of cell types in 3D co-cultures. Reducing CellTracker Green to 0.5X improved JE6.1 cell visibility and minimized over-labelling artifacts. (E) ATP-based CTG viability assay for co-cultures stained with combinations of CellTracker and SPY650 nuclear dye showed that the optimized staining cocktail (CellTracker Orange 2X + CellTracker Green 0.5X + SPY650) did not compromise viability, supporting its compatibility with live-cell assays. (F) Quantification of segmented cells per condition revealed that 0.5X CellTracker Green significantly increased the number of JE6.1 cells detected within optimal size and intensity range, while staining with 2X CellTracker Orange preserved segmentation efficiency for 786-O cells. *Scale bars: 50 µm.* Error bars indicate mean ± SEM; *p < 0.05, **p < 0.01.

The optimized staining conditions significantly enhanced the quality of downstream 3D segmentation and quantification. The combination of SPY650 (24 hour incubation) and CellTrackers (2X/0.5X, 45min incubation prior culturing), cultured for 72 h, provided distinct labelling of cell nuclei and cytoplasm, facilitating robust single-cell segmentation and accurate quantification in both monoculture and co-culture spheroids. These optimized protocols are well-suited for HT 3D single-cell analysis and enable more detailed characterization of drug responses in complex cell models (Supp. Table 1, Supp. Fig. 1F). We additionally evaluated the optimized staining protocol in ccRCC patient-derived cancer cells (PDCs) co-cultured with peripheral blood mononuclear cells (PBMCs) under previously described culture conditions^29^ (see Methods). SPY650 enabled clear nuclear visualization, while differential CellTracker labeling allowed the cancer and immune cell populations to be distinguished (Supp. Fig. 1G). CTG measurements showed no detectable reduction in metabolic activity under the optimized staining conditions (Supp. Fig. 1H), providing proof-of-concept support for application of the staining strategy to more sensitive patient-derived models.

**Table 1.** Components and criteria of the objective function for evaluating 3D single-cell segmentation quality.

| Objective function component | Description | Criteria |
| --- | --- | --- |
| **Cell number and size range** | Measures the total number of segmented cells. Only cells having a volume within a realistic biological size range are acceptable. | Cell volumes must fall within a biologically plausible operational range (900–8200 $\mu\text{m}^3$; see Methods). Penalization occurs if the number of cells within this range is too low (explained below). |
| **Pixel coverage** | Maximizes pixel coverage to capture entire cells rather than just nuclei, ensuring correct cell boundary identification without over-segmentation. | Calculates the proportion of image pixels that fall within the final watershed masks, relative to the total number of pixels in the image, to assess segmentation coverage. |
| **Cell morphology (cell extent)** | Measures the extent of segmented cells as a proxy for irregular or fragmented cell shapes. | Extent is calculated as the ratio of segmented object volume to bounding-box volume. The objective function uses 1-extent as a shape penalty, assigning higher penalties to objects with lower extent, as may occur with elongated, irregular, or fragmented segmentations. |
| **Cell number penalty** | Penalizes segmentations with too few valid cells based on biological size expectations. Encourages a sufficient number of realistically sized segmented cells. | A penalty is applied when fewer than $x$ cells fall within the valid volume range (~900–8200 $\mu\text{m}^3$). The penalty decreases linearly as the count approaches $x$, with a maximum penalty of 0.5 when no valid cells are detected, and no penalty when $x$ or more are present. In our specific case, $x$ is 150 cells. |

### Image preprocessing and pretrained 3D U-Net deep learning models achieve single-cell segmentation of dense 3D spheroids

To improve input images for 3D DL-based segmentation, we evaluated channel merging, intensity scaling, and contrast enhancement. The previously optimized CellTracker Orange cytoplasmic and SPY650 nuclear channels were merged after scaling the cytoplasmic signal, with mean merging and scaling factors between 5 and 20 producing balanced visualization of cell boundaries and internal structures (Fig. 4A). We subsequently compared histogram equalization, Contrast Limited Adaptive Histogram Equalization (CLAHE), and contrast stretching, with CLAHE^30^ and contrast stretching producing the most uniform and well-defined single-cell profiles (Fig. 4B). These observations were used to define plausible parameter ranges for subsequent Bayesian optimization.

**Figure 4.**
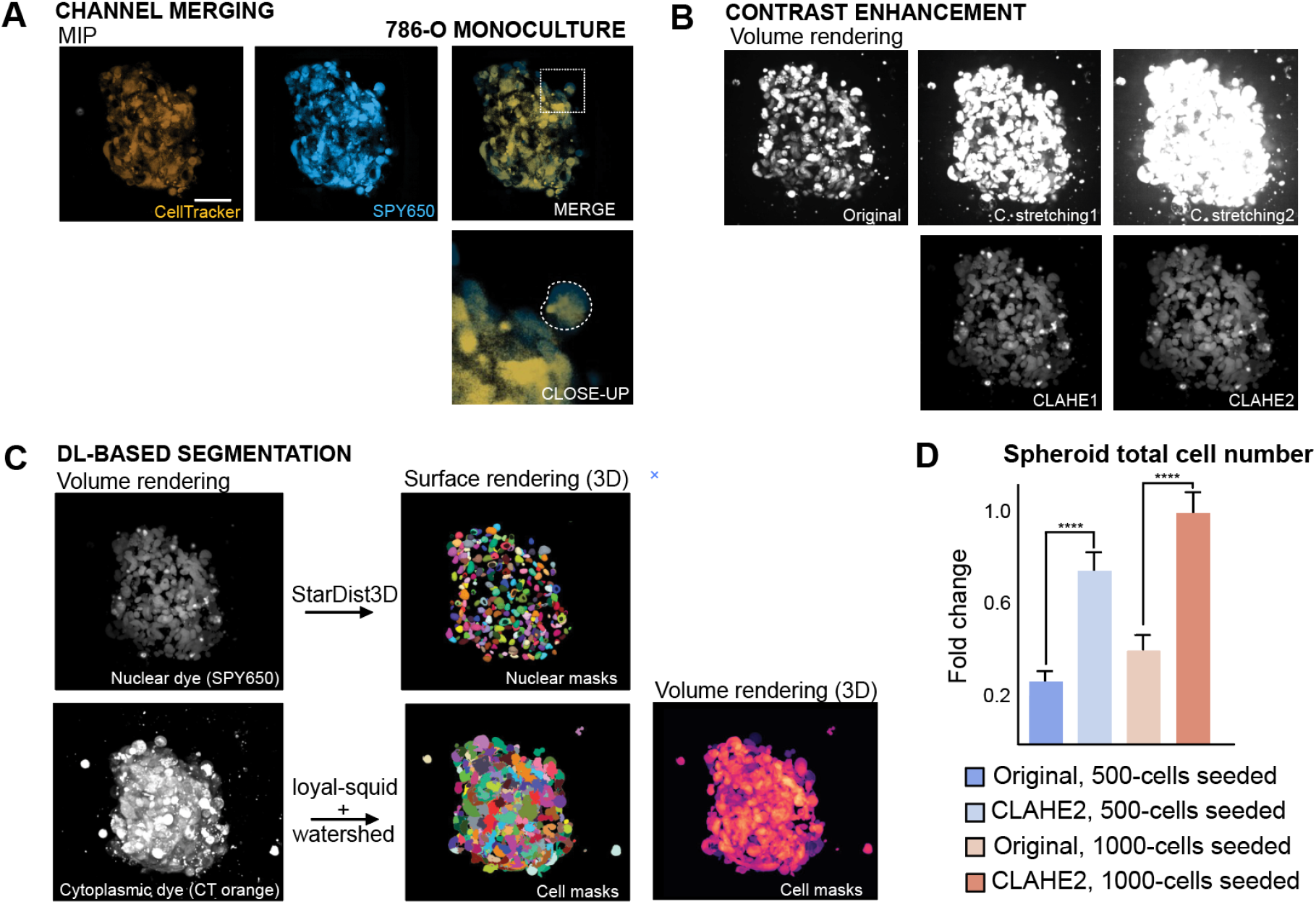
Image preprocessing and DL models for single-cell segmentation of dense 3D spheroids. 786-O 3D cultures were used for image processing parameter evaluation. (A) Cells were stained with CellTracker and SPY650 for cytoplasmic and nuclear detection; the merging was done using the channels’ means after scaling the cytoplasmic signal by a factor of 10, producing a balanced composite image. This step is referred to as brightness adjustment in the code and involves scaling through pixel intensity multiplication. The close-up image highlights a single cell (dashed circle) where both cytoplasmic and nuclear signals are visible, offering a more complete visualization of individual cell morphology. (B) Contrast enhancement methods—CLAHE and contrast stretching—were used and evaluated to improve cell boundary differentiation. (C) DL-based segmentation pipeline using nuclear (SPY650) and cytoplasmic (CellTracker Orange) channels as inputs. StarDist3D was used for nuclear segmentation, while whole-cell masks were generated using the loyal-squid 3D U-Net model with watershed refinement. Volume-rendered input images are shown on the left. The center panels display 3D surface renderings of nuclear and cell masks in pseudo-color, highlighting individual objects. The right panel shows a 3D volume rendering of the final cell masks, providing an overview of the spheroid’s structure and cell distribution. (D) Total segmented cell number, expressed as fold change, for spheroids seeded with 500 or 1000 cells and segmented from original or CLAHE2-enhanced images. CLAHE2 enhancement resulted in higher segmented cell counts at both seeding densities. Error bars indicate mean ± SEM; ****p < 0.0001.

Segmenting densely packed and heterogeneous cells within 3D spheroids presents a major challenge, particularly in the absence of reliable ground truth annotations. This challenge is further amplified in high-content (HC) confocal screening datasets, where large numbers of spheroids must be analyzed automatically^22^. Manual annotation is therefore impractical for such complex datasets, necessitating alternative segmentation strategies. Our pipeline combined deep learning models for nuclear and whole-cell segmentation with additional image-processing methods to address this challenge. For nuclear segmentation we used the StarDist3D model^31^, which successfully generated precise nuclear masks even in areas where cells were tightly clustered, leveraging its pre-trained capacity for complex 3D nuclear segmentation (Fig. 4C). This model enabled effective initial segmentation, providing foundational markers for the subsequent cell segmentation steps.

For whole-cell segmentation, we utilized the loyal-squid 3D U-net model^32^, originally developed using embryonic mouse tissue imaged with light-sheet microscopy^33^. Preliminary testing of image resizing and voxel interpolation did not improve segmentation performance; therefore, the original image dimensions were retained for the final segmentation pipeline. The combination of StarDist3D-derived nuclear masks and loyal-squid whole-cell masks allowed for the effective implementation of watershed segmentation to further refine cell boundaries (Fig. 4C). This multi-step approach, integrating DL and classical segmentation methods, provided robust and reliable segmentation results across various conditions. The refined segmentation pipeline enabled accurate extraction of key features, such as cell count, volume, and texture. Comparisons between different contrast enhancement methods (e.g., original versus CLAHE-enhanced images) demonstrated that image preprocessing markedly affected segmentation results, including the number of detected cells (Fig. 4D), motivating the subsequent Bayesian optimization of preprocessing parameters.

### Bayesian optimization provides robust parameter selection for enhanced segmentation quality

Visual inspection of segmentation results provided an initial understanding of the parameter space and helped establish reasonable ranges for distinguishing individual cells in 3D spheroids. However, the subjective nature of visual assessment highlighted the need for a more quantitative and systematic approach for optimizing the segmentation pipeline. To address this, we implemented Bayesian optimization to automatically identify optimal parameter combinations for image preprocessing and segmentation while defining an objective function that served as a surrogate for ground truth (see Supp. text A & B).

The Bayesian optimization was applied using a Gaussian process–based minimization approach. This approach allowed us to efficiently explore the high-dimensional parameter space of the segmentation pipeline, which included parameters for image preprocessing steps (e.g., brightness adjustment, contrast enhancement) and deep-learning models (e.g., tiling size, foreground thresholds). The optimization process aimed to minimize an objective function that quantitatively evaluated segmentation quality based on several predefined criteria. Key components of the objective function are included in Table 1.

The general formula of the objective function, *O*, is given by

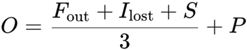

where

● *F_out_* is the mean of the frequency (ratio) of objects outside the desired size range.
● *I_lost_* is the mean percentage of image pixels not covered by the final segmentation masks.
● *S* is the mean extent-based shape penalty (1 − extent), where extent is the ratio of segmented object volume to bounding-box volume; lower S values indicate objects occupying a greater proportion of their bounding box.
● *P* is the mean of the cell count penalty that is added if the number of cells segmented falls below a certain threshold.

The parameter space included a range of preprocessing and segmentation parameters such as brightness scaling, contrast enhancement settings, and DL model thresholds, which were defined as either categorical or continuous variables. Details of all optimized parameters are provided in the Methods section (Supp. Table 2, and Supp. text A). Bayesian optimization was used to efficiently search for parameter combinations that minimized the objective function.

**Table 2.** Drug panel and experimental concentrations.

| Name | Mechanism/<br>Targets | Class | Supplier<br>Ref | Supplier | Concentration (nM) |
| --- | --- | --- | --- | --- | --- |
| Epacadostat | IDO inhibitor | Immunomodulator | HY-1568<br>9 | Medchem<br>Express | 10-1000-10<br>000 |
| Lenalidomide | Immunomodulatory | Immunomodulator | HY-A000<br>3 | Medchem<br>Express | 100-10000<br>- 100000 |
| Dactolisib | mTOR/(PI3K)<br>inhibitor | Kinase inhibitor | HY-5067<br>3 | Medchem<br>Express | 1-100-1000 |
| Temsirolimus | binds FKBP12,<br>causes inhibition<br>of mTORC1 | Rapalog | T-8040 | LC<br>Laboratories | 0.1-10-100 |
| Pirfenidone | Antifibrotic and<br>anti-inflammatory | Immunomodulator | CT-PIRF | ChemieTek | 10-1000-10<br>000 |

### Objective function values reflect segmentation quality

We first evaluated the ability of our custom objective function to guide segmentation optimization using an initial Bayesian optimization run comprising 10 objective-function evaluations. The optimization outcomes were strongly associated with improvements in segmentation quality: lower objective function values corresponded to well-defined cell boundaries, minimal fragmentation, and reduced overlap between segmented objects. In contrast, segmentations with higher objective function values produced visually poorer segmentation outcomes, including merged objects and fragmented segmentations (Fig. 5A). The pipeline converged on a parameter set that reduced the objective function value by approximately 56% relative to the baseline (Fig. 5B). These observations confirm that the objective function provides a reliable surrogate for visual segmentation quality in the absence of ground truth annotations. To extend this validation, we performed an extended Bayesian optimization run comprising 80 objective-function evaluations. This experiment demonstrated consistent convergence of the objective function across successive evaluations (Fig. 5C). In parallel, we monitored each component of the objective function (Supp. Fig. 2). Across the optimization, nearly all components showed progressive improvements, supporting the internal consistency and interpretability of the objective function as a multidimensional quality readout.

**Figure 5.**
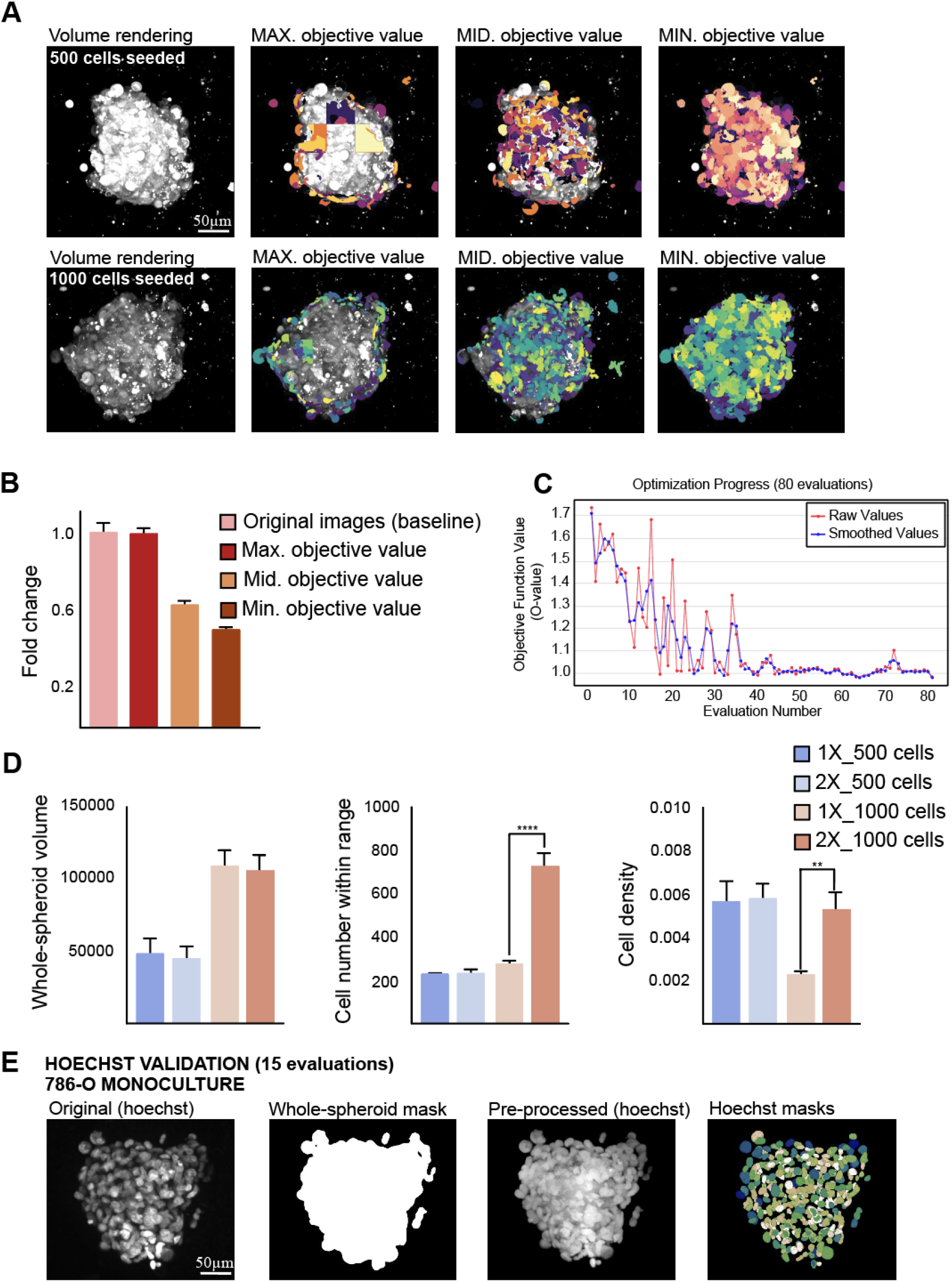
Bayesian optimization enhances segmentation performance in 3D spheroids. (A) Representative segmentation results corresponding to high, intermediate, and low objective function values obtained during Bayesian optimization of 786-O spheroids seeded (500 or 1000 cells seeded). Lower objective function values correspond to improved segmentation quality, with well-separated and clearly delineated individual cells (individual segmentation masks are shown in different colors). (B) Objective function values relative to the baseline parameter set. The optimized parameter set reduced the objective function by approximately 56%. (C) Optimization progress across 80 objective-function evaluations, showing the raw objective-function values (red) and a rolling mean calculated over three consecutive evaluations (blue), illustrating the convergence trend. (D) Quantitative comparison of segmentation outputs under different CellTracker Orange concentrations (1X and 2X) and seeding densities (500 and 1000 cells). Spheroids seeded with 1000 cells and stained with 2X CellTracker yielded the highest number of segmented cells within the expected size range and the greatest measured cell density (**P < 0.01; ****P < 0.0001). (E) Validation of pipeline transferability using Hoechst-stained 786-O monocultures. A Bayesian optimization run comprising 15 objective-function evaluations generated high-quality segmentation masks and reduced the objective function by approximately 50%, despite differences in nuclear dye and image characteristics. *Scale bars: 50 µm.* Error bars indicate mean ± SEM.

We then applied the optimized pipeline to segment spheroids seeded with different initial densities and stained with varying CellTracker Orange concentrations. Notably, spheroids seeded with 1000 cells and stained with 2X CellTracker yielded the highest number of segmented cells within the expected size range and the greatest measured cell density compared with the other conditions (Fig. 5D). These results demonstrate that the optimized pipeline can be successfully applied across biologically distinct experimental conditions, while segmentation outputs appropriately reflect differences in staining intensity and initial cell seeding.

Finally, to assess the generalizability of our approach across different imaging conditions, we performed a separate Bayesian optimization run comprising 15 objective-function evaluations on Hoechst-stained 786-O spheroids. Despite the different nuclear dye and image texture, the pipeline achieved an approximately 50% reduction in the objective function and generated high-quality segmentation outputs (Fig. 5E). This result highlights the adaptability of our Bayesian optimization framework to diverse staining modalities and supports its utility for HT segmentation across heterogeneously labelled 3D datasets.

### Validation of MorphoNavigator-3D for automated cell-type quantification in 3D co-cultures

To evaluate the accuracy of our 3D image-based segmentation pipeline for quantifying single cells in spheroid cultures, we first compared segmentation-derived counts with flow cytometry measurements across monoculture (786-O cancer cells) and co-culture (786-O + JE6.1 T-cells) conditions (Fig. 6). Following 3D imaging, the same spheroids were dissociated into single-cell suspensions and analyzed by flow cytometry. CellTracker Orange and CellTracker Green fluorescence enabled discrimination between cancer cells and T-cells, allowing direct comparison of absolute cell counts and cell-type proportions obtained by image-based segmentation and flow cytometry. Both methods showed consistent relative differences between experimental conditions, although segmentation systematically yielded lower cell counts than flow cytometry. This difference reached statistical significance in the monoculture condition (p < 0.05; Fig. 6A). To explain this discrepancy, we generated a schematic of the U-bottom well system used in our assays (Fig. 6B). This discrepancy is likely explained by the imaging field being centered on the spheroid, thereby excluding some free-floating cells that were subsequently detected by flow cytometry. Bland–Altman analysis revealed a mean difference of −178 cells (± SD: 195 cells), with most observations falling within the 95% limits of agreement (Fig. 6C). In addition, segmentation-derived cell counts showed a strong linear relationship with flow cytometry measurements (R² = 0.94; Fig. 6D), indicating that MoNa-3D captures relative differences in cell abundance across experimental conditions while preserving spatial information that is inherently lost during flow cytometric analysis. Segmentation also yielded similar relative proportions of cancer and T cells in co-cultures compared with flow cytometry (Fig. 6E). T-cell counts were more variable, likely due to their non-adherent behavior, as the segmentation approach focuses on the spheroid and may exclude peripheral or loosely attached immune cells.

**Figure 6.**
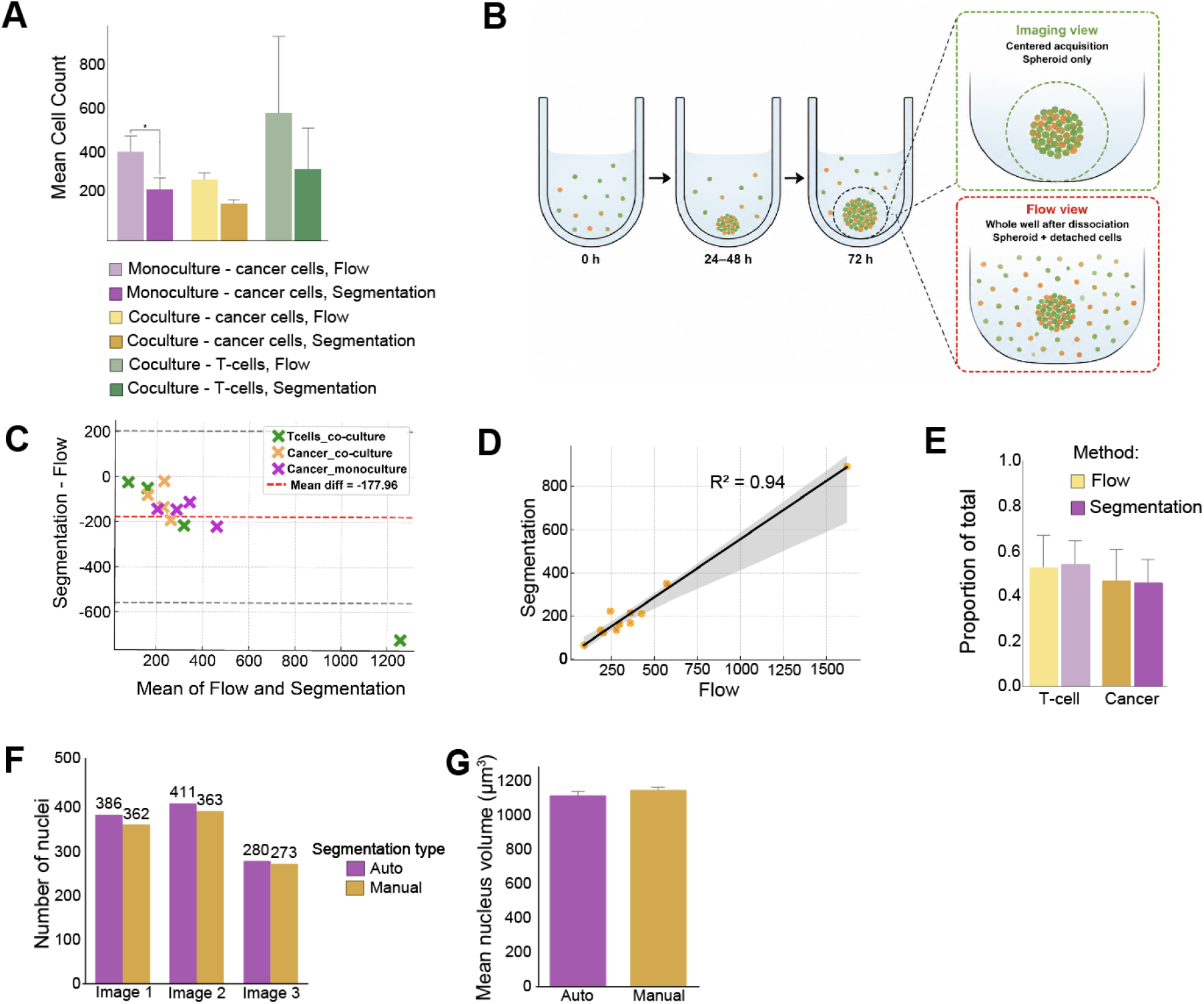
Comparison of image-based segmentation with flow cytometry and manual segmentation for cell quantification in 3D cultures. (A) Mean cell counts obtained from flow cytometry and image segmentation in monoculture and co-culture conditions. Error bars represent SEM. A statistically significant difference was observed in monocultured cancer cells (*p < 0.05). (B) Schematic illustration of the U-bottom well culture system showing spheroid formation and the distribution of floating and aggregated cells over time. The diagram highlights that image segmentation captures primarily the spheroid and its immediate surroundings, whereas flow cytometry captures all cells present in the well, including those outside the imaging field. (C) Bland–Altman plot assessing agreement between segmentation-derived and flow cytometry cell counts. The red dashed line indicates the mean difference between methods (−177.96 cells), with gray dashed lines marking the 95% limits of agreement. (D) Linear relationship between segmentation-derived and flow cytometry cell counts (R² = 0.94). (E) Cell-type proportions in co-culture (T-cells vs. cancer cells) as estimated by flow cytometry and segmentation. Despite underestimation of total cell numbers, segmentation provided consistent compositional ratios between cell types. (F) Comparison of the number of nuclei identified by the automated pipeline versus manual segmentation in monocultures, demonstrating consistency between the two methods (n = 3). (G) Bar plot showing no statistical difference in mean nucleus volume between automated and manual segmentation (n = 3, p = 0.6260). Error bars indicate mean ± SEM; *p < 0.05, **p < 0.01.

To benchmark MoNa-3D pipeline against manual annotation, a subset of spheroids was manually segmented in 3D using napari^34^, and cell counts and nuclear volumes were compared between the two approaches. MoNa-3D-derived cell counts were highly similar to manually annotated counts (7–24 nuclei difference, n = 3; Fig. 6F) and mean nuclear volumes did not differ significantly (paired t-test, n = 3, P = 0.6260; Fig. 6G). Minor differences in the spread of nuclear volumes (mean ± SD: pipeline 1111 ± 785 μm³, manual 1136 ± 535 μm³, n = 3) likely reflect variability in how low signal nuclei were identified manually, and the more cylindrical 3D objects produced manually in napari, which could reduce variability in nuclear shape measurements relative to the more rounded StarDist3D reconstructions. Notably, manual segmentation required careful annotation lasting up to 10 hours per image, whereas MoNa-3D completed analysis of a single image in approximately 2 min on a high-performance computer (described in the Methods).

Together, these results demonstrate that MoNa-3D provides rapid and reliable quantitative measurements that closely agree with manual annotation, supporting its use for HT phenotypic profiling and comparative studies in 3D co-culture systems.

### Phenotypic profiling of drug-induced effects in 3D co-cultures using single-cell segmentation and spatial analysis

We applied MoNa-3D (Supp. Text B) to investigate phenotypic responses of ccRCC (786-O) and T-cell (JE6.1) co-cultures to five drugs with distinct mechanisms of action: Dactolisib and Temsirolimus (PI3K/mTOR pathway inhibitors), Pirfenidone, Lenalidomide, and Epacadostat (immunomodulatory compounds)^35–38^ (Fig. 7A). Bayesian optimization was extended to 80 objective-function evaluations across the drug-treated images, with convergence of the objective-function values indicating stabilization of the optimization process (Supp. Fig. 4A). This enabled the integration of ATP-based viability with morphological, compositional, regional, and spatial measurements (Supp. Table 3)^39–45^. Dactolisib and Temsirolimus produced the strongest reductions in ATP-based viability and spheroid volume, whereas Pirfenidone, Lenalidomide, and Epacadostat showed comparatively modest effects (Fig. 7B,C).

**Figure 7.**
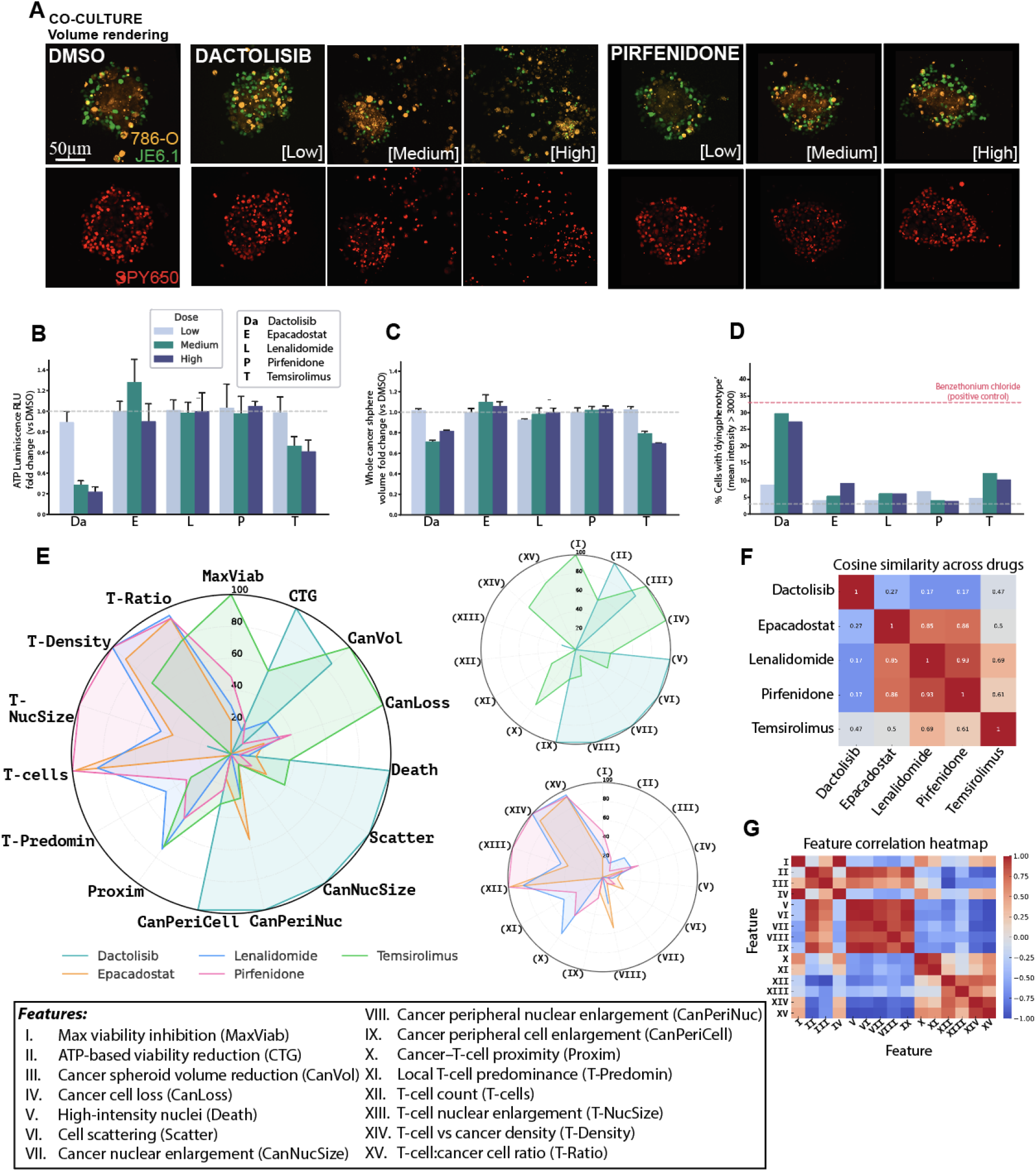
Application of the optimized pipeline: multidimensional phenotypic profiling of drug responses in cancer/T-cell 3D co-cultures. (A) Overview of the experimental setup: 3D volume renderings of co-cultures composed of 786-O renal cancer cells (CellTracker Orange) and JE6.1 T-cells (CellTracker Green), co-stained with SPY650 nuclear dye. Representative images are shown for DMSO control, Dactolisib, and Pirfenidone treatments across three drug concentrations (low, medium, high). (B) Quantification of ATP-based cell viability by CTG luminescence assay, represented as fold-change relative to DMSO control (shown dotted line) across increasing drug concentrations. Dactolisib and Temsirolimus displayed pronounced reductions in ATP levels consistent with ATP-based viability loss, while immunomodulatory compounds induced minor to moderate changes. (C) Whole-spheroid volume measured from reconstructions of the cancer cell fluorescence channel (orange), shown as fold-change compared to DMSO control (shown dotted line). (D) Fraction of cancer cells presenting high nuclear intensities (mean voxel intensity >3000), used as a proxy for nuclear condensation and cell death, especially upon Dactolisib treatment. (E) Global phenotypic radar (circular) maps displaying 15 multidimensional feature profiles across treatments after min-max scaling and directionality correction. Each axis corresponds to a distinct phenotypic feature, integrating viability, spheroid architecture, cellular composition, nuclear morphology, regional remodeling, and tumor–immune spatial organization parameters. Drug-specific phenotypic fingerprints show distinct response patterns for PI3K/mTOR inhibitors (Dactolisib and Temsirolimus; upper radar plot) compared with the immunomodulatory compounds (Epacadostat, Lenalidomide, and Pirfenidone; lower radar plot). (F) Pairwise cosine similarity between normalized drug-level phenotypic profiles. Higher values indicate greater similarity between multidimensional response profiles. (G) Pearson correlation matrix across the 15 phenotypic features, illustrating coordinated relationships among viability, morphology, composition, and spatial measurements. *Scale bars: 50 µm.* Error bars indicate mean ± SEM.

Each readout was selected to capture a distinct biological dimension of drug response. Metabolic activity was quantified as the mean luminescence signal per well using the CellTiter-Glo (CTG) ATP-based viability assay, a well-established pharmacological endpoint^38^. Imaging-derived measurements comprised spheroid volume and cell scattering as surrogates of tissue architecture disintegration^40,41^, nuclear enlargement as an indicator of drug-induced polyploidy or senescence (often associated with mTOR inhibition)^42,43^, and chromatin condensation intensity as a proxy for cell death^44^. In addition, population-specific spatial metrics were used to quantify cancer–T-cell proximity, local cross-population neighbourhood structure, and relative cell composition^19–21,38,45^.

The PI3K/mTOR inhibitors also produced cell-type-specific effects, with both Dactolisib and Temsirolimus reducing T-cell counts, whereas the other compounds produced comparatively modest changes in segmented cancer-and T-cell numbers (Supp. Fig. 3A,B). Under Dactolisib, cancer-cell counts remained relatively preserved despite the pronounced reduction in ATP-based viability and, at the highest concentration, were slightly higher than in DMSO controls, although still below the initial seeding density of 500 cells (Fig. 7B; Supp. Fig. 3A). Thus, the number of morphologically detectable cancer cells did not directly reflect the bulk metabolic response. Dactolisib also increased cancer-cell nuclear enlargement, particularly at the spheroid periphery (Supp. Fig. 3C; Supp. Fig. 4D), and markedly increased high-intensity cancer-cell nuclei (Fig. 7D). These phenotypes are consistent with cellular states associated with growth arrest, senescence, polyploidy, mTOR inhibition, and cellular stress or cell death^42–44,46^; high-intensity nuclei were also prominent in the cytotoxic positive control.

Spatial analyses further revealed treatment-dependent remodeling not captured by cell counts alone. Dactolisib and Temsirolimus increased overall cell scattering, consistent with spheroid disorganization (Supp. Fig. 3E). Dactolisib produced the greatest cancer–T-cell separation and highest cancer: T-cell ratio, together with increased cancer-cell dispersion relative to local T-cell coverage (Supp. Fig. 3F,G). Temsirolimus showed an intermediate spatial phenotype, with increased cancer-cell dispersion but comparatively preserved local cancer–T-cell association, whereas Pirfenidone, Lenalidomide, and Epacadostat generally preserved local cross-population organization (Supp. Fig. 3F–H). Together, these measurements distinguished global structural dispersion from local tumor–immune spatial remodeling^19,20,45^.

Integration of 15 viability, morphological, compositional, regional, and spatial features generated distinct multidimensional response profiles across treatments (Supp. Table 3; Fig. 7E). Dactolisib showed the strongest overall phenotype, characterized by ATP-based viability and spheroid-volume reduction, nuclear remodeling, reduced T-cell abundance, and disrupted tumor–immune spatial organization. Temsirolimus shared several viability and structural effects but comparatively preserved local cancer–T-cell association. In contrast, Epacadostat, Lenalidomide, and Pirfenidone showed more limited structural disruption and greater preservation of T-cell abundance and spatial organization. These integrated profiles therefore captured both tumor-intrinsic responses and immune-associated spatial changes that were not apparent from bulk viability measurements alone.

Cosine similarity confirmed Dactolisib as the most divergent integrated response, while Epacadostat, Lenalidomide, and Pirfenidone were more similar and Temsirolimus occupied an intermediate position (Fig. 7F). Feature correlation identified coordinated relationships among viability, nuclear remodeling, cell scattering, and cellular composition (Fig. 7G). Complementary single-cell and treatment-level analyses similarly distinguished the two cell populations and treatment-associated phenotypes, with regional analysis showing pronounced peripheral cancer-cell enlargement following Dactolisib and Temsirolimus treatment (Supp. Fig. 4B–E). Together, the multidimensional profiles distinguished the pronounced structural, cellular, and spatial effects of PI3K/mTOR inhibition from the comparatively preserved spheroid organization observed with the other compounds, while also resolving differences between Dactolisib and Temsirolimus.

## DISCUSSION

MorphoNavigator-3D is a high-throughput framework for multidimensional phenotypic profiling of 3D spheroids at single-cell resolution. Conventional 3D segmentation workflows often rely on manual annotations or labor-intensive ground-truth datasets that are impractical for densely packed multicellular spheroids^10^. By integrating optimized live-cell staining, DL-based segmentation, and Bayesian optimization guided by an interpretable objective function, MoNa-3D enables automated, annotation-free optimization across experimental conditions, complementing emerging integrated 3D high-content screening platforms such as HCS-3DX42^22^.

Compatibility with live-cell dyes (SPY650 and CellTracker) facilitates integration with endpoint assays such as ATP-based viability and flow cytometry^15,16^. Segmentation was maintained under challenging drug-induced conditions, including spheroid disaggregation and nuclear remodeling, while comparison with flow cytometry showed consistent estimates of relative cell composition while preserving spheroid architecture and spatial information. Application of MoNa-3D to ccRCC–T-cell co-cultures exposed to drugs with distinct mechanisms revealed phenotypic responses not captured by ATP-based viability alone. The panel included PI3K/mTOR pathway inhibitors and immunomodulatory compounds relevant to targeted therapy and immune modulation^47–51^.

Single-cell measurements revealed distinct responses to Dactolisib and Temsirolimus despite reductions in ATP-based viability and spheroid volume with both treatments. Dactolisib induced pronounced structural disintegration, T-cell loss, cancer-cell dispersion, reduced local cancer–T-cell association, and cancer-cell nuclear remodeling. Notably, cancer-cell counts remained relatively preserved despite reduced ATP-based viability, but remained below the initial seeding density in both Dactolisib-treated and control spheroids and therefore should not be interpreted as evidence of continued proliferation. Instead, morphologically detectable cancer cells persisted alongside nuclear enlargement and high-intensity nuclear phenotypes consistent with altered cellular states including growth arrest, senescence, polyploidy, severe stress, or cell death^42,44,52^. Because CTG measures the combined metabolic activity of both cell populations, their relative contributions to the reduced ATP signal cannot be resolved from the bulk assay alone. Instead, the cell-type-resolved imaging measurements revealed distinct responses of the two populations, with pronounced morphological remodeling of the cancer-cell compartment occurring alongside loss of T-cell representation. Temsirolimus similarly reduced ATP-based viability and T-cell abundance but produced less extensive cancer-cell nuclear remodeling and comparatively preserved local cancer–T-cell association. These differences may reflect selective mTORC1 inhibition by Temsirolimus versus dual PI3K/mTOR inhibition by Dactolisib, although their mechanistic basis was not directly tested^48,50,53^. The diverging phenotypes illustrate how MoNa-3D can distinguish cell-type-specific morphological, compositional, and spatial responses that are obscured in bulk measurements. Together, the observed intensity, morphological, compositional, and spatial phenotypes are consistent with previous reports linking PI3K/mTOR inhibition to nuclear remodeling, cell-cycle arrest, and cell death^48,50,53^.

Regional analysis revealed preferential nuclear and cytoplasmic enlargement at the spheroid periphery following Dactolisib and Temsirolimus treatment, potentially reflecting spatial differences in drug exposure or cellular state^54^. This spatial heterogeneity parallels observations in ccRCC tissues, where tumor states and tumor–immune architectures vary across anatomical regions and have been associated with therapeutic response^55,56^. Multiplex spatial tissue profiling has likewise identified aggressive epithelial–mesenchymal transition (EMT)-associated and immunosuppressive phenotypes at the invasive tumor border^57^. In comparison, Pirfenidone, Lenalidomide, and Epacadostat generally preserved spheroid architecture, tumor–immune organization, and T-cell representation to a greater extent than Dactolisib^5,18^. By resolving drug responses across spheroid center, middle, and periphery, MoNa-3D provides an in vitro analogue of this spatial organization, enabling quantitative characterization of how local remodeling and immune interactions shape global phenotypic states. Together, these findings suggest that regional spatial metrics may serve as informative readouts for evaluating immunomodulatory responses *in vitro* and underscore the translational importance of spatial characterization for predicting therapeutic responsiveness.

Multivariate analysis further distinguished drug classes and resolved intra-class differences between Dactolisib and Temsirolimus, particularly in tumor–immune organization, peripheral morphology, and cancer-cell dispersion relative to T-cell neighborhoods. The broader phenotype associated with Dactolisib is consistent with the more extensive effects reported for dual PI3K/mTOR inhibition^48,58^. Quantification of multidimensional phenotypes may inform precision oncology strategies and support rational drug combination design by capturing structural remodeling and immune organization that are not resolved by conventional viability assays. Distinct phenotypic dimensions affected by Dactolisib and the non-PI3K/mTOR compounds may also help prioritize mechanistically complementary drug combinations for subsequent experimental testing. This aligns with growing efforts to improve the physiological relevance of preclinical drug-testing models by incorporating multicellular complexity and immune interactions^59^. Finally, the phenotypic maps generated by MoNa-3D provide a standardized and visually interpretable framework for comparing multidimensional drug responses across experimental systems, including patient-derived organoids.

Several limitations remain. The annotation-free objective function requires broader validation across staining conditions, cell types, and imaging platforms, and segmentation-derived cell counts reflect morphologically detectable cells rather than their functional or metabolic state. Moreover, JE6.1 cells provide a standardized but immortalized, non-antigen-specific T-cell model; consequently, the spatial measurements reported here should not be interpreted as direct measures of antigen-specific immune activity. Although evaluated only in a single proof-of-concept ccRCC patient-derived cancer cell–PBMC co-culture, the optimized staining protocol did not detectably reduce metabolic activity and enabled discrimination between cancer and immune cell populations, providing preliminary evidence for its applicability to patient-derived ex vivo models. Validation across additional patient-derived and autologous tumor–immune systems, together with extension to multiplexed phenotypic assays such as Cell Painting, will therefore be important for establishing broader applicability^15,60^.

Altogether, MorphoNavigator-3D provides an adaptable framework for multidimensional single-cell phenotyping in multicellular 3D systems, integrating automated segmentation with morphological and spatial analysis to characterize drug responses beyond bulk viability measurements.

## MATERIALS AND METHODS

### 3D cultures

Monocultures of the clear cell renal cell carcinoma cell line 786-O (CRL-1932; ATCC) were cultured for 3 days to form 3D spheroids. For tumor–immune co-culture experiments, 786-O cells were combined with the T-cell line Jurkat Clone E6-1 (JE6.1; TIB-152; ATCC). HUVEC endothelial cells (CRL-1730; ATCC) were additionally used in selected experiments during optimization and evaluation of the staining and imaging workflow. Cells were maintained in RPMI-1640 supplemented with 10% FBS, 100 U/mL penicillin, and 100 µg/mL streptomycin, and incubated at 37°C with 5% CO₂. Monocultures were seeded at 500 cells/well unless otherwise specified. Co-cultures were seeded at 500 cells/well (786-O) and 250 cells/well (JE6.1). Spheroids were grown in ultra-low attachment 384-well round-bottom plates (Corning, CLS3830) under scaffold-free conditions. Patient-derived cancer cells (PDCs) from a clinically diagnosed ccRCC patient were isolated as previously described^28,60^ and cultured in a 3:1 (v/v) mixture of Ham’s F-12 Nutrient Mix (11765054; Gibco) and DMEM (11965092; Gibco). The PDC model used in this study was derived from a high-risk ccRCC tumor and has been previously genetically characterized by whole-exome sequencing, which confirmed the preservation of clinically relevant somatic variants between the parental tumor and the derived PDC culture, including a VHL mutation characteristic of ccRCC^29,62^. The medium was supplemented with 5% FBS, 8.4 ng/mL cholera toxin (Sigma, C8052), 0.4 µg/mL hydrocortisone (Sigma, H4001), 10 ng/mL epidermal growth factor (Corning, 354052), 24 µg/mL adenine (Sigma, A8626), 5 µg/mL insulin (Sigma, 91077C), and 10 µM Y-27632 Rho kinase inhibitor (Sigma, SCM075). Culture surfaces were coated with 2% Matrigel prior to seeding. Cell line authentication for 786-O and JE6.1 was performed at the Institute for Molecular Medicine Finland (FIMM) Genomics Unit (HiLIFE, University of Helsinki).

### Fluorescent labelling

For far-red nuclear labeling, either NucSpot (1µM for 1X, 40082-T, biotium), SYTO (1µM for 1X, S34900; Invitrogen), or SPY650-DNA (1 µM for 1X, Spirochrome AG) were prepared according to the manufacturer’s instructions and added to the spheroid cultures at different timepoints as outlined in the Results section. For cytoplasmic labeling, cells were first labelled with the CellTracker dyes (at 0.5 µM, 1 µM, or 2µM) (C2925 and C34551; Thermo Fisher Scientific) for 45min, before seeding them into the wells for 3D culturing. The labelled cells were allowed to aggregate and form spheroids over a period of 3 days. Other cytoplasmic dyes included PKH26 (2µM or 2X, PKH26GL; Sigma-Aldrich.), CellVue (2µM or 2X., MINCLARET; Sigma-Aldrich) with a similar incubation approach.

### Viability assays

**a. ATP-based cell viability assessment (CTG Assay)**: Cell viability was measured using the luminescence-based CellTiter-Glo (CTG) assay (Promega), which measures ATP levels as a proxy for the number of metabolically active (viable) cells. Luminescence readings were obtained using a PHERAstar FS plate reader (BMG Labtech). Spheroids were treated with CTG reagent according to the manufacturer’s protocol, and luminescence was measured from the 384-well plates directly.
**b. LIVE/DEAD assay:** A LIVE/DEAD dual marker assay (R37601; Invitrogen) was conducted to evaluate the level of apoptosis and cell death under the optimized dye conditions. Spheroids were stained with calcein-AM and ethidium homodimer-1 and imaged to quantify live (green) and dead (red) cells, for 1 hr before imaging, under the manufacturer’s instructions.
**c. DRAQ7**: DRAQ7 (D15105; Invitrogen), a far-red fluorescent DNA dye, was used to assess cell membrane permeability and cell death in spheroids. Spheroids were incubated with DRAQ7 (final concentration, 3 µM) for 1 h at 37 °C in a humidified incubator with 5% CO₂ before imaging. The stained spheroids were then imaged, and the percentage of DRAQ7-positive cells was quantified to evaluate cell death under the experimental conditions.

### Flow cytometry-based cell quantification

Flow cytometry was used to validate the number and proportion of single cells in co-cultures. Spheroids were dissociated into single-cell suspensions by adding 50 μL of 0.25% Trypsin-EDTA directly to the wells and incubating at 37°C for 5–10 min intervals, with gentle pipetting after each cycle to aid in dissociation without vortexing. When approximately 95% of cells appeared dissociated as singlets (typically within 45 min), fresh culture medium was added to each well to reach a final volume of 95 μL. The cell suspensions were then transferred to conical-bottom 96-well plates (Nunc) for flow cytometry. Flow cytometry-based cell profiling was performed by the FIMM High Throughput Biomedicine Unit (HTB, HiLIFE, University of Helsinki). Dead cells were excluded using DRAQ7 (Cell Signaling Technologies, #7406), added directly to the wells and incubated for 10 min at room temperature. Single cells were analyzed on iQue Plus and iQue3 HT flow cytometers (Sartorius). Data were analyzed using ForeCyt software (Sartorius). Gating was performed to exclude DRAQ7-positive dead cells and identify CellTracker Orange-and CellTracker Green-positive populations corresponding to cancer cells and T-cells, respectively. Absolute cell counts were extracted per population for comparison with image-based quantification.

### Drug Sensitivity and Resistance Testing (DSRT)

Drug sensitivity and resistance testing (DSRT) was conducted using 3D spheroid cultures of the renal cancer cell line 786-O, as well as co-cultures with T-cells (JE6.1). The drugs were dissolved in dimethyl sulfoxide (DMSO) or water and dispensed into 384-well plates at three increasing concentrations over a 10,000-fold concentration range (Table 2). Spheroids were cultured in ultra-low attachment 384-well round-bottom plates (Corning, CLS3830) containing the indicated drugs or 0.1% DMSO vehicle control. Cells were labeled with CellTracker dyes, seeded in 25 μl volume of RPMI medium and incubated for 72 h at 37°C with 5% CO₂ with drugs. After 48 h, nuclei were stained with SPY650. After 72 h of drug incubation and imaging, cell viability was measured using the luminescence-based CellTiter-Glo (CTG) assay (Promega) to assess the efficacy of each drug. Luminescence counts were obtained using a PHERAstar FS plate reader (BMG Labtech).

### Manual segmentation

A subset of SPY650-stained spheroids was manually segmented by a single expert annotator using the built-in labeling tool in napari and the *napari-label-interpolator* plugin which connects annotations of top, middle and bottom slice of each nucleus^63^. The resulting annotations were used to compare segmentation metrics with those obtained from the automated MoNa-3D pipeline. For both manual and automated segmentation, only nuclei within a whole spheroid mask, generated using the same tools, were included in the comparison analyses, as detached nuclei were difficult to identify reliably by eye. Nuclei counts and volumes were extracted for comparison.

### Image acquisition

Images were acquired using an Opera Phenix high-content spinning-disk confocal microscope (Revvity; FIMM High-Content Imaging and Analysis core unit; FIMM-HCA, HiLIFE, University of Helsinki, Finland) equipped with a 20× water immersion objective (NA 1.0) and 405, 488, 561, and 640 nm lasers. Image stacks were acquired in confocal mode using laser autofocus and comprised 45 optical sections. The resulting 3D images had a voxel size of 0.299 × 0.299 × 2.5 µm (x, y, z) and an image size of 2160 × 2160 pixels per optical section. Images were initially saved in the native Opera Phenix format and subsequently converted to 3D TIFF format for downstream analysis.

### Computational analysis and optimization

Signal intensity, dye penetration depth, and spheroid integrity were assessed to determine optimal staining conditions. Images were initially analyzed using Harmony software (Revvity) and BIAS (Single-Cell Technologies Ltd.), whereas napari was used for visualization of segmentation outputs. The segmentation pipeline and Bayesian optimization framework were implemented as custom Python scripts executed in Jupyter notebooks using NumPy, pandas, OpenCV, scikit-image, tifffile, StarDist, bioimageio.core, SciPy, and scikit-optimize (skopt). The scripts performed image preprocessing (e.g., channel merging and contrast enhancement), whole-spheroid and single-cell segmentation using deep-learning models, and Bayesian optimization of segmentation parameters. Routine analyses were performed on a workstation equipped with an AMD Ryzen 9 7900X 12-Core processor (4.70 GHz), 64 GB RAM, and an NVIDIA GeForce RTX 4070 Ti GPU (12 GB VRAM), whereas computationally intensive Bayesian optimization runs were performed on the FIMM High-Performance Computing GPU cluster (Institute for Molecular Medicine Finland, HiLIFE, University of Helsinki) using a server including an NVIDIA A100 40 GB GPU, 256 GB RAM, and half of the AMD EPYC 7402 24-core CPU resources.

#### a. Segmentation parameter optimization

Bayesian optimization was used to explore a multidimensional space of image-preprocessing and segmentation parameters by minimizing a composite objective function based on segmentation quality metrics (Table 1). Optimized variables included image intensity scaling, adaptive histogram equalization (CLAHE), contrast stretching, and the watershed foreground threshold. Parameter ranges and fixed inference settings are summarized in Supplementary Table 2.

All categorical and continuous optimization parameters were defined using the scikit-optimize (skopt) library and optimized using Gaussian process regression with the expected improvement acquisition function. Optimization was performed over multiple iterations, with the objective function guiding the search toward parameter combinations that maximized segmentation quality. Tiling and halo sizes were treated as inference configuration parameters rather than optimization variables. They were selected during preliminary testing to balance GPU memory requirements and minimize tile-edge artifacts and were fixed for the final optimization runs (tile_xy = 256, tile_z = 10, halo_xy = 32, halo_z = 2). These parameters remain user-configurable in the released software to facilitate adaptation to different image dimensions, model architectures, and hardware configurations. During preliminary testing, certain tiling and halo combinations produced inference errors due to dimensional mismatches or insufficient overlap. These configurations were automatically identified and excluded using validation logic (Supplementary Text A).

The quality of each candidate segmentation was evaluated using a composite objective function (Table 1) combining biologically and morphologically motivated criteria. One component assessed the proportion of segmented objects falling within a biologically plausible cell-volume interval of 900–8200 µm³. This interval was selected as a permissive, dataset-specific range informed by experimentally measured mammalian cell volumes across multiple cultured cell lines^64–66^, representative tumor-cell volumes reported in computational models of tumor growth^67^, and empirical inspection of correctly segmented cells in the present datasets. The selected limits were intended to preserve the expected biological variability of single-cell volumes within the experimental system while excluding fragmented objects and likely merged segmentations. Cell volumes were calculated by converting segmented voxel counts into physical units using the calibrated image voxel dimensions (0.299 × 0.299 × 2.5 µm).

#### Regional analysis

To investigate spatially resolved drug effects, segmented single cells were assigned to one of three spheroid regions—core, middle, or periphery—based on their 3D Euclidean distance from the spheroid centroid. The centroid was calculated from the whole-spheroid mask. Cells were then binned into tertiles according to their distance: the core included the inner 0–33rd percentile of distances, the middle covered the 34th–66th percentile, and the periphery included the outermost 67th–100th percentile. This distance-based stratification enabled regional analysis of nuclear and cytoplasmic features, cell densities, and drug-induced phenotypic alterations.

#### Spatial and neighborhood analysis

Spatial measurements were calculated from the 3D centroids of segmented cancer cells and T-cells after conversion of voxel coordinates to physical distances using the calibrated image dimensions. Global cell scattering was quantified from intercellular Euclidean distances between segmented cell centroids. Cancer–T-cell proximity was quantified using cross-population distance measurements, with lower distances indicating closer spatial association between the two populations. To characterize local tumor–immune organization beyond global distance measurements, population-specific neighborhood analyses were performed using a 20 µm radius around each segmented cell. This radius was selected to capture a short-range multicellular neighborhood corresponding approximately to one to two mammalian cell diameters while minimizing contributions from more distant cells within the spheroid. For each cancer cell, the number of neighboring T-cells located within 20 µm was quantified; conversely, for each T-cell, the number of neighboring cancer cells within the same radius was calculated. These local cross-population neighborhood measurements were combined with population-specific dispersion metrics to derive cancer-centric and T-cell-centric dispersion-to-neighbor ratios. Higher ratios therefore reflect greater spatial dispersion relative to local cross-population coverage, whereas lower ratios indicate comparatively preserved local cellular association.

#### Multivariate feature extraction and normalization

Following segmentation, a comprehensive set of quantitative single-cell features describing cell morphology, geometry, intensity, and spatial properties was extracted for each segmented object. From these measurements, biologically relevant viability, morphological, compositional, and spatial descriptors were derived, and a representative subset of 15 features (Supp. Table 3) was selected for multivariate analysis. These selected features were normalized across all drug conditions using min–max scaling (range 0–1) to enable direct comparisons. Where necessary, feature directionality was inverted so that higher normalized values consistently reflected stronger drug-induced phenotypic effects. Directionally transformed features included cancer-cell loss, cell scattering, ATP-based viability inhibition, and immune infiltration metrics.

Feature correlation, phenotypic similarity, and dimensionality reduction: Pearson correlation coefficients were calculated across the selected phenotypic features to identify coordinated and potentially redundant measurements. Feature pairs with |R| ≥ 0.85 were flagged as strongly correlated. Pairwise cosine similarity was calculated between normalized drug-level feature vectors to quantify similarity between integrated phenotypic response profiles. Radar plots were used to visualize the multidimensional profiles, while principal component analysis and hierarchical clustering were used as complementary unsupervised approaches to assess treatment-level phenotypic structure.

#### Statistics

ATP-based viability, cell counts, spheroid volume, and segmentation scores were compared across groups using mean ± SEM. Data distributions were assessed for normality (Shapiro-Wilk) and variance homogeneity (Levene’s test). Depending on distribution, either one-way ANOVA or Kruskal-Wallis tests were applied. Pairwise comparisons to DMSO control were performed using Mann-Whitney U tests with Bonferroni correction. Mean nuclear volumes from manual versus automated segmentation were compared across experiments using a paired t-test.

#### Code availability

The code used for image preprocessing, whole-spheroid and single-cell segmentation, and Bayesian optimization is available in GitHub: https://github.com/bioimage-profiling/Bayesian-segmentation3D

#### Data availability

The datasets generated and analyzed during this study will be deposited in Zenodo and made publicly available upon publication. The accession link and DOI will be provided before final publication.

## Acknowledgements

We thank Katja Välimäki and Antti Hassinen (FIMM, HiLIFE, University of Helsinki) for their excellent technical assistance. We are grateful to Constantin Pape and Anwai Archit for their invaluable guidance at the start of the segmentation pipeline development. We also thank the FIMM High-Throughput Biomedicine Unit (FIMM-HTB) for preparation of pre-drugged assay plates, and designing and performing the flow cytometry experiments, as well as the FIMM High Content Imaging and Analysis unit (FIMM-HCA; Euro-BioImaging) for their imaging services and expertise and FIMM Genomics. All aforementioned FIMM units are supported by HiLIFE and Biocenter Finland. Figures were created using BioRender.com, Inkscape and Adobe Illustrator. We acknowledge the iCAN Flagship Project of the Research Council of Finland, the Finnish Cancer Center -FICAN, the Cancer Foundation of Finland, the Research Council of Finland (340273, 346604, 359907), and Finnish Cultural Foundation-Kymenlaakso Foundation, for grants enabling this research. P.H acknowledge support from TKP2021-EGA09 (TKP-31-8/PALY-2021), TKCS-2024/73, Horizon-BIALYMPH, Horizon-SYMMETRY, Horizon-RIA SWEEPICS (101135053), Horizon 2020 FAIR-CHARM (101016457), HAS-NAP3 (NAP2022-I-6/2022), HUNRENTECH (TECH-2024-34), OTKA-SNN (139455/ARRS), and NKKP (ADVANCED_24_151202).

## Author contributions

I.M., M.F., V.P., and L.P. conceived the study. I.M., M.F., V.P., and L.P. developed the methodology. I.M. developed the MoNa-3D computational pipeline and Bayesian optimization framework, with contributions from P.N. and G.A. I.M. conducted the computational analyses, formal analysis, data curation, and generated the visualizations. I.M. and M.F. designed the biological experiments. I.M., M.F., A.N., and A.M. performed the biological experiments. I.M., M.F., and A.N. analyzed the biological data. A.N. generated the manual segmentation annotations and contributed to figure preparation. P.N. contributed to the conceptual development and implementation of the Bayesian optimization framework and objective function. G.A. contributed to computational infrastructure, GPU implementation, code optimization, and preparation of the GitHub repository. A.R. facilitated access to the patient-derived samples used in this study through the DEDUCER study. V.C. provided the PBMC samples used for the patient-derived co-culture experiments. P.H., V.P., and L.P. supervised the study. P.H., V.P., and L.P. acquired funding. I.M., V.P., A.N., and L.P. wrote the original manuscript. All authors reviewed, edited, and approved the final version of the manuscript.

## Ethical issues

Written informed consent was obtained from the patient. The study was conducted under protocols approved by the Ethics Committee of the Hospital District of Helsinki and Uusimaa (HUS) (Dnro 154/13/03/02/2016, approved 15 March 2017; HUS/850/2017, approved 14 July 2021). Patient samples and corresponding clinical data were collected as part of the DEDUCER study under an Institutional Ethical Review Board-approved protocol and in accordance with the principles of the Declaration of Helsinki. The use of blood samples from healthy donors complied with the requirements of the Finnish Red Cross Blood Service (26/2025). All study procedures were performed in accordance with applicable local legislation and institutional requirements, and the annotation and handling of personal data followed the principles of the General Data Protection Regulation (GDPR).

## Competing interests

P.H. is affiliated with Single-Cell Technologies Ltd. The remaining authors declare no competing interests.

## Notes

https://github.com/bioimage-profiling/Bayesian-segmentation3D

### Competing Interest Statement

P.H. is affi liated with Single-Cell Technologies Ltd. The remaining authors declare no competing interests.

### Summary of Updates

This revised version substantially expands the original study and introduces MorphoNavigator-3D, an integrated framework for automated, annotation-free single-cell analysis of 3D cancer spheroids. The manuscript now includes a schematic overview of the complete experimental and computational workflow, an expanded and optimized live-cell staining strategy, and further development of the Bayesian optimization framework for deep-learning-based 3D segmentation. We added extensive validation of the image-analysis pipeline, including comparison of segmentation-derived cell counts and cell-type proportions with flow cytometry, Bland-Altman and regression analyses, and benchmarking against manual 3D segmentation. The drug-response study was expanded from two compounds to a five-drug panel spanning PI3K/mTOR inhibitors and immunomodulatory compounds, together with additional single-cell morphological, regional, compositional, and cancer-T-cell spatial analyses. These measurements were integrated into multidimensional phenotypic maps to compare drug-response profiles and resolve both inter- and intra-class phenotypic differences. We additionally demonstrate application of the optimized staining strategy to a patient-derived ccRCC-PBMC co-culture. The manuscript, figures, supplementary information, author list, Methods, and Discussion have been extensively revised and expanded accordingly.

