## Supplementary Material for "MorphoNavigator-3D: Generalizable single-cell phenotyping of cancer spheroids using Bayesian-optimized deep-learning workflows"

### Supp. MATERIAL

**Supp. Table 1.** optimized fluorescence live dyes conditions for single-cell visualisation of spheroids.

| Dye | Purpose | Concentration | Incubation time | Signal intensity | Notes |
|---|---|---|---|---|---|
| **SPY650** | Nuclear staining (far-red) | 1X | 24 h | Bright | Best signal depth and clarity in 3D spheroids. 72 h incubation causes spheroid degradation. |
| **CellTracker** | Cytoplasmic staining (blue-red-orange-green) | 2X (for 786-O cells), 0.5X (for JE6.1 cells) | 72 h | Bright | Uniform staining, consistent signal across cell types. 0.5X concentration increases immune cell visibility. |

**Supp. Table 2.** Parameters used for Bayesian optimization.

| Parameter | Role | Type | Search range / Final value | Applied to |
|---|---|---|---|---|
| **Brightness adjustment factor** | Optimized | Categorical | 0, 1, 10, 15, 20 | Cytoplasmic channel (before merging) |
| **CLAHE clip limit** | Optimized | Real | 0.0–0.5 | Per channel |
| **CLAHE number of bins** | Optimized | Categorical | 128, 256, 444, 512 | Per channel |
| **Contrast stretching lower percentile** | Optimized | Integer | 1–20 | Per channel |
| **Contrast stretching upper percentile** | Optimized | Integer | 90–99 | Per channel |
| **Contrast stretching scale range** | Optimized | Categorical | e.g. [0.5,1.2], [0.8,1.3] | Per channel |
| **Foreground threshold** | Optimized | Real | 0.2–0.7 | Watershed input |

| | | | | |
|---|---|---|---|---|
| **Tiling size** | Fixed | Categorical | tile_xy = 256; tile_z = 10 | Loyal-squid inference |
| **Halo size** | Fixed | Categorical | halo_xy = 32; halo_z = 2 | Tile overlap |

*Note:* Tiling and halo parameters were optimized during preliminary method development and fixed for the final Bayesian optimization reported in this study. The released software retains these parameters as user-configurable settings to facilitate adaptation to different datasets and hardware configurations.

**Supp. Table 3.** Phenotypic features integrated into the multidimensional drug-response profiles.

| Feature | Biological interpretation | Metric definition |
|---|---|---|
| **Maximum viability inhibition (MaxViab)** | Maximum treatment-associated reduction in ATP-based viability response | Maximum inhibition value derived from the drug-response measurement across tested concentrations |
| **ATP-based viability reduction (CTG)** | Reduction in metabolic activity/ATP content | CTG luminescence normalized to DMSO and transformed so higher values indicate greater ATP reduction |
| **Cancer spheroid volume reduction (CanVol)** | Loss of spheroid structural volume | Whole-spheroid volume derived from 3D cancer fluorescence segmentation and normalized to DMSO |
| **Cancer-cell loss (CanLoss)** | Reduction in segmented cancer-cell abundance | Number of segmented cancer cells within the accepted volume range, normalized to DMSO and directionally transformed |
| **High-intensity nuclei (Death)** | Nuclear condensation/stress-associated phenotype | Fraction of cancer-cell nuclei exceeding the predefined mean nuclear-intensity threshold (>3000) |
| **Cell scattering (Scatter)** | Loss of compact multicellular organization | Mean 3D intercellular distance between segmented cell centroids |
| **Cancer nuclear enlargement (CanNucSize)** | Drug-associated nuclear remodeling | Proportion of cancer cells with nuclear volume >4000 $\mu m^3$ |

| | | |
|---|---|---|
| **Cancer peripheral nuclear enlargement (CanPeriNuc)** | Region-specific nuclear remodeling | Relative increase in cancer-cell nuclear size within the spheroid periphery compared with DMSO |
| **Cancer peripheral cell enlargement (CanPeriCell)** | Region-specific whole-cell remodeling | Relative increase in cancer-cell size within the spheroid periphery compared with DMSO |
| **Cancer–T-cell proximity (Proxim)** | Spatial association between tumor and immune populations | Cross-population cancer–T-cell distance metric, transformed so higher plotted values indicate closer association |
| **Local T-cell predominance (T-Predomin)** | Local representation of T-cells around cancer cells | Population-specific local neighborhood score derived from the number of T-cell neighbors around cancer cells within a 20 μm radius |
| **T-cell count (T-cells)** | T-cell abundance | Number of segmented T-cells within the accepted volume range, normalized to DMSO |
| **T-cell nuclear enlargement (T-NucSize)** | T-cell nuclear remodeling | Proportion of T-cells with nuclear volume >4000 $\mu m^3$ |
| **T-cell vs cancer density (T-Density)** | Relative population density within the spheroid | Ratio/relative comparison of T-cell and cancer-cell densities, with each population normalized to measured whole-spheroid volume per cell type |
| **T-cell:cancer-cell ratio (T-Ratio)** | Relative cell-population composition | Ratio of segmented T-cells to segmented cancer cells within the accepted cell-volume range |

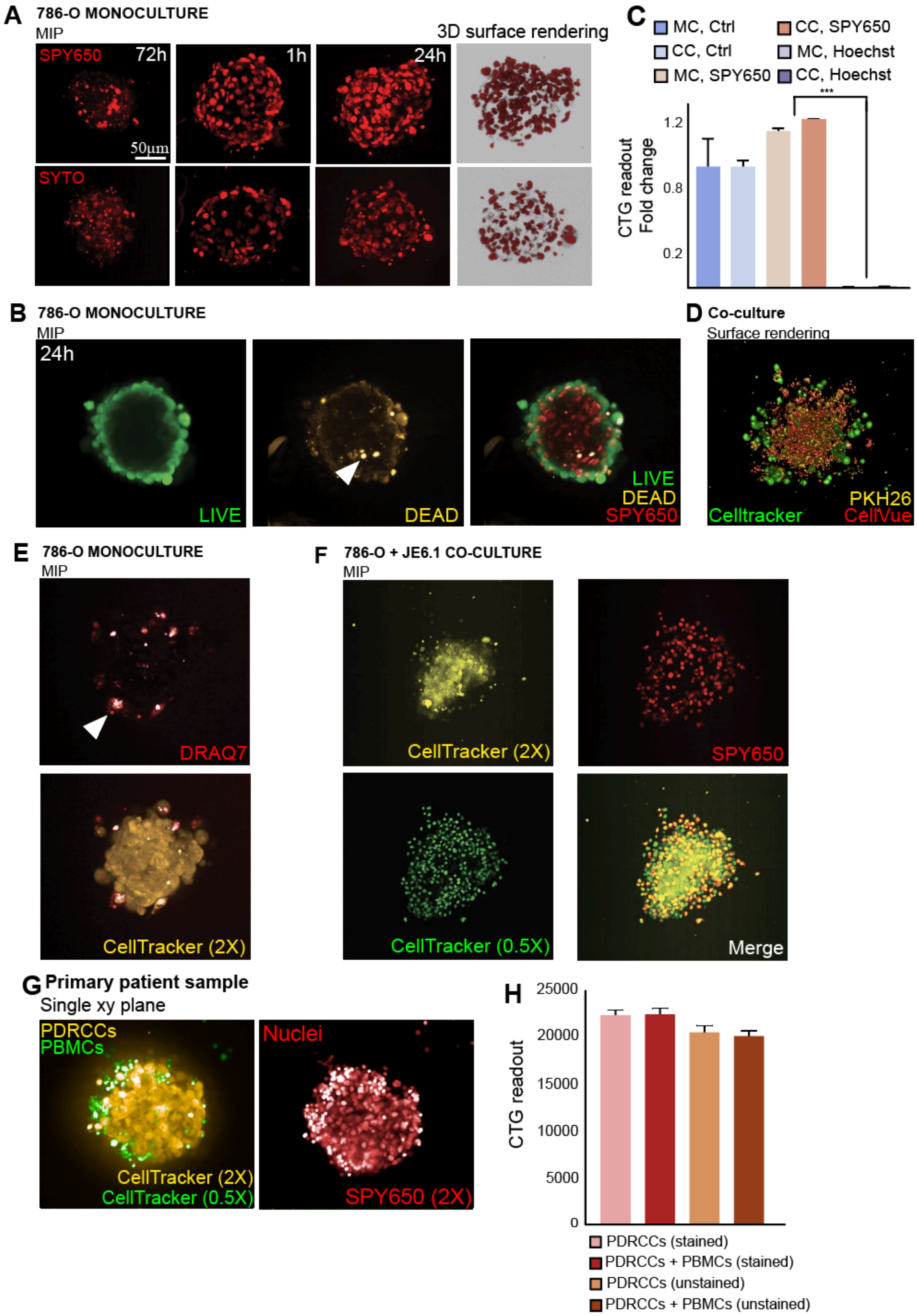


**Supplementary Figure 1. Evaluation of fluorescence dyes' incubation times, concentrations and toxicity in 3D spheroids.** (A) Time-course evaluation of nuclear staining in 786-O spheroids using SPY650 and SYTO dyes. MIPs and 3D surface renderings are shown for SPY650 (top row) and SYTO (bottom row) at 72 h, 1 h, and 24 h post-incubation. SPY650 produced a robust nuclear signal after 24 h, while 72 h incubation resulted in spheroid disintegration, indicating potential toxicity. SYTO showed similar patterns as SPY650 but with lower signal intensity. (B) LIVE/DEAD dual marker assay (LIVE: green; DEAD: yellow) confirms low cell death in 786-O spheroids after 24 h SPY650 incubation. The merged image includes SPY650 staining (red), highlighting colocalization with apoptotic cells. (C) CTG ATP-based viability assay showing fold change in metabolic activity under different staining conditions in mono- (MC) and co-cultures (CC). SPY650 did not significantly affect viability, whereas Hoechst staining resulted in a strong reduction in signal, particularly in co-culture (***$p < 0.001$). (D) Co-culture of T-cells (JE6.1, CellTracker green), cancer (786-O, PKH26 red) and endothelial cells (HUVEC, CellVue yellow); CellTracker produced the most consistent and uniform cytoplasmic staining unlike PKH26 and CellVue, which exhibited uneven, punctuated patterns. (E) Validation of CellTracker Orange (2 µM) and DRAQ7 (dead cell marker) in 786-O spheroids. CellTracker provided uniform labelling, while DRAQ7 signal was restricted to a few cells (arrowhead), supporting minimal toxicity. (F) Optimized staining of 786-O + JE6.1 co-cultures using CellTracker Green (0.5X), CellTracker Orange (2X), and SPY650. The merged image shows distinct labelling of the two populations and successful nuclear visualization. (G) Optimized staining of patient-derived renal cancer cells (PDRCCs; CellTracker Orange, 2X) and PBMCs (CellTracker Green, 0.5X) with SPY650 nuclear dye (2X) enables clear population distinction. (H) ATP-based CTG viability measurements show no significant differences between stained and unstained conditions, confirming minimal dye toxicity (G). *Scale bars*: 50 *µm*. Error bars indicate mean ± SEM.

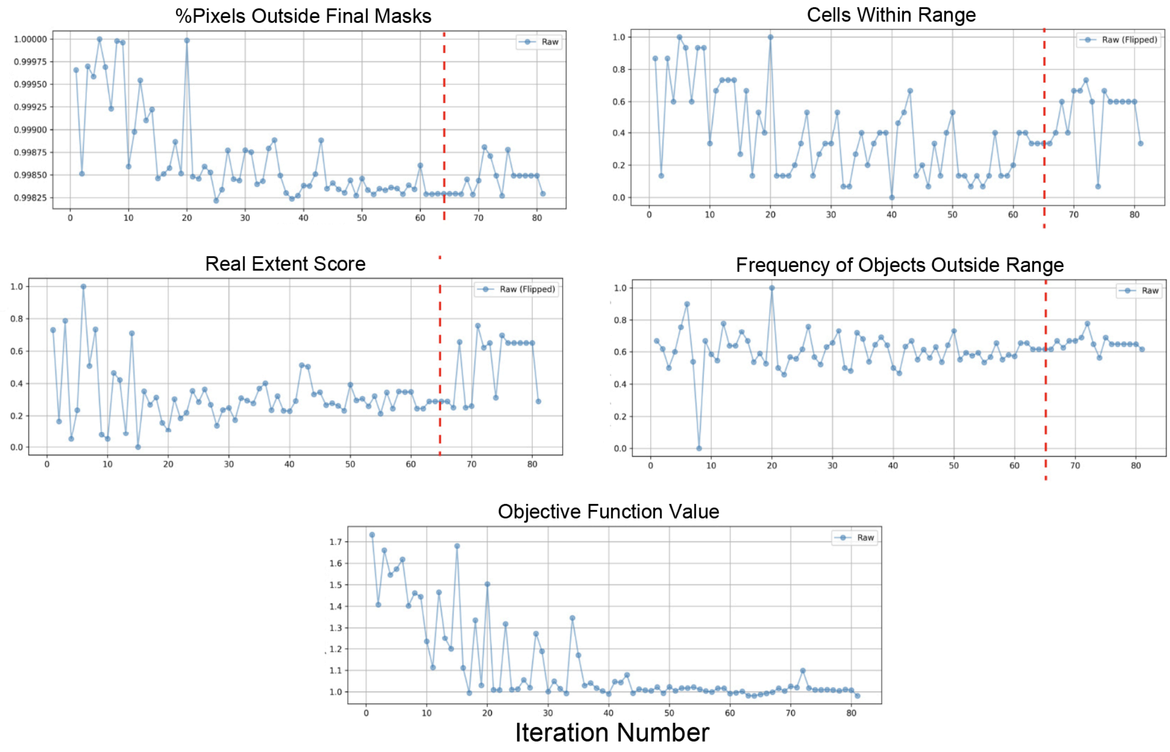


**Supplementary Figure 2. Multi-metric breakdown of segmentation optimization progress.** Tracking of individual metrics used to define the composite objective function throughout 80 iterations of Bayesian optimization in 3D co-cultured spheroids: Top left: Percentage of pixels falling outside final masks, which decreased steadily as segmentation improved. Top right: Percentage of segmented cells falling within the biologically expected size range (flipped score for minimization). Middle left: Extent score of real segmented cells (flipped score for minimization), increasing over time as segmentation became cleaner. Middle right: Frequency of segmented objects falling outside the expected feature range. Bottom: Composite objective function value summarizing all metrics; converging toward a minimum by iteration 60. The red dashed line indicates the start of parameter flipping to refine convergence. These plots present the robustness and convergence behavior of the objective-driven optimization approach.

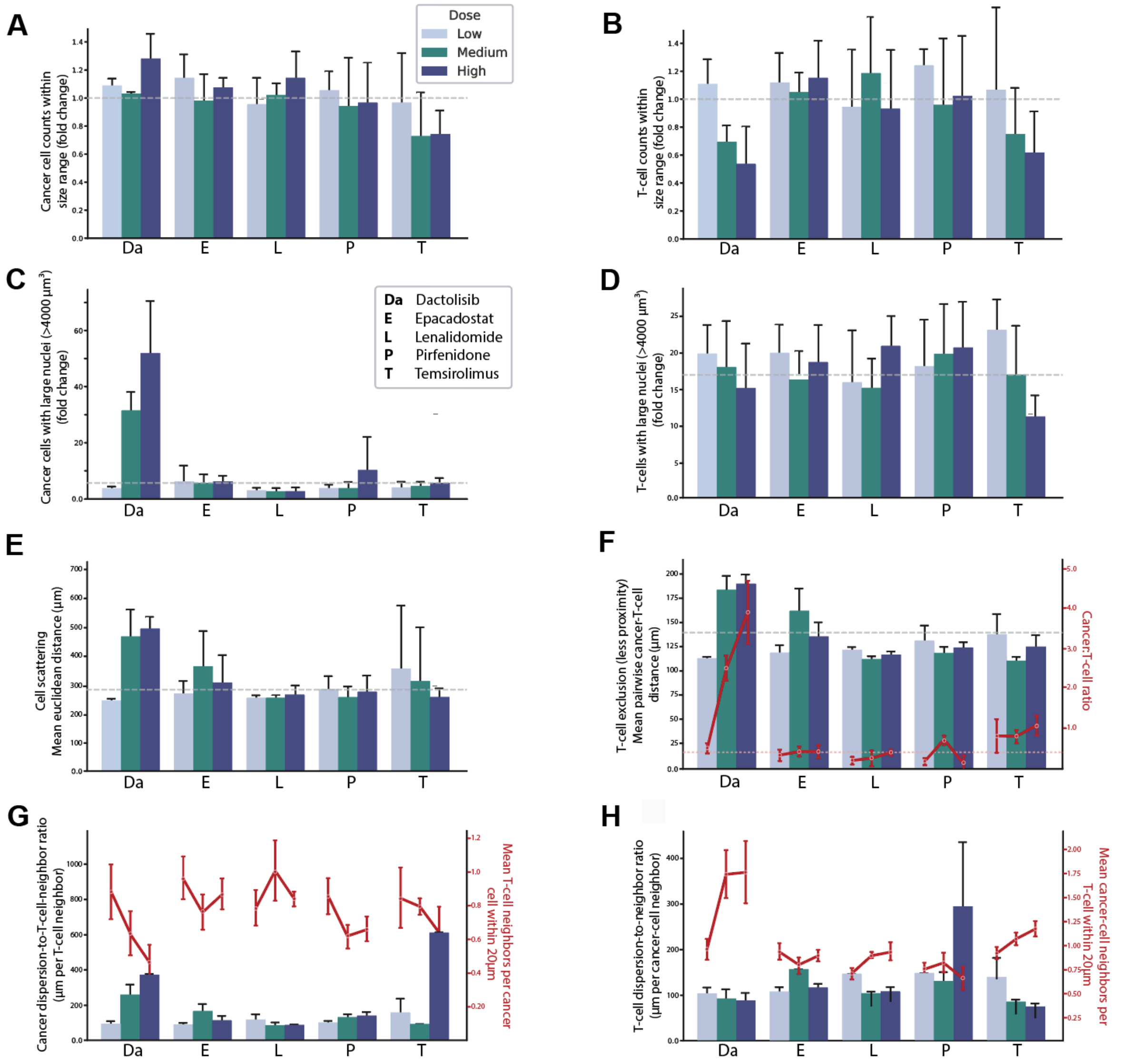


**Supplementary Figure 3. Additional cell count and nuclear morphology analyses across drug treatments.** (A,B) Fold change in the number of segmented cancer cells (A) and T-cells (B) within the accepted cell volume range (900–8200 µm³), normalized to the DMSO control. (C,D) Fold change in the number of segmented cancer cells (C) and T-cells (D) with enlarged nuclei (nuclear volume >4000 µm³), normalized to the DMSO control. Bars represent the low, medium, and high drug concentrations. The dashed line indicates the DMSO control (fold change = 1). Error bars represent mean ± SEM. (E) Mean Euclidean distance between segmented cell centroids, used as a measure of overall cell scattering. (F) Cancer–T-cell spatial proximity across treatment conditions, with larger distances indicating reduced local association. Bars show the cross-population cancer–T-cell distance metric, while the red line shows the corresponding cancer:T-cell count ratio, providing compositional context for the spatial measurement. (G) Cancer-centric spatial organization quantified as cancer-cell dispersion relative to local T-cell coverage. Bars show the dispersion-to-neighbor ratio, and the red line indicates the mean number of T-cell neighbors per cancer cell within a 20 µm radius. (H) T-cell-centric spatial organization quantified as T-cell dispersion relative to local cancer-cell context. Bars show the corresponding dispersion-to-neighbor ratio, and the red line indicates the mean number of cancer-cell neighbors per T-cell within a 20 µm radius. Bars represent low, medium, and high drug concentrations. Dashed lines indicate the corresponding DMSO control values. Error bars represent mean ± SEM.

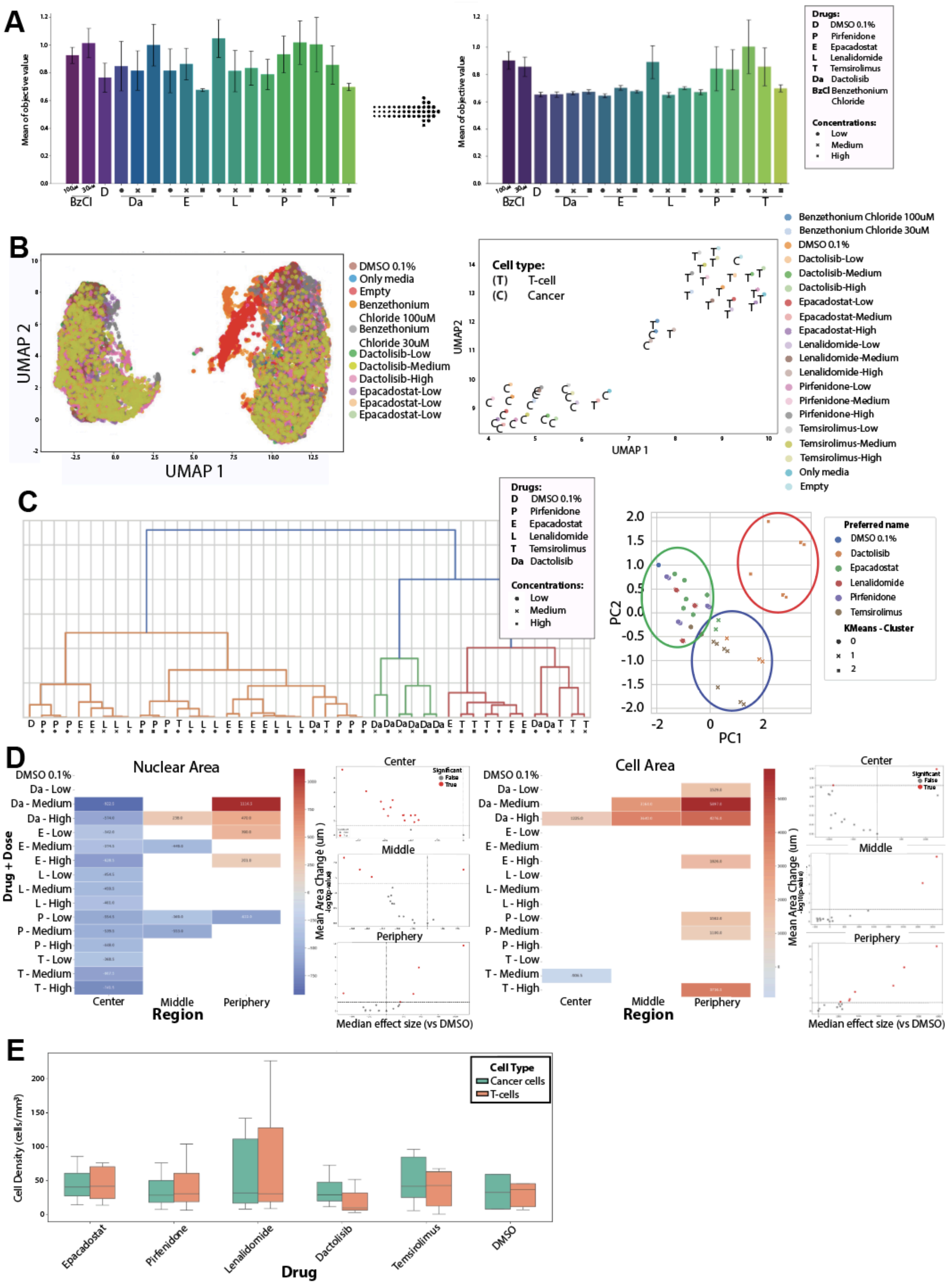


**Supplementary Figure 4. Regional and multivariate analysis of drug-induced phenotypes in 3D co-cultures using single-cell image features and integrated drug response metrics.** (A) Objective function values obtained after 15 (left) and 80 (right) cycles of Bayesian optimization across drug conditions. Extended optimization (80 cycles) resulted in more stable and consistent segmentation outcomes across treatments, reflected by reduced variance in objective scores. (B) UMAP projections of segmented single-cell morphological features. Left: projection of individual cells colored by the drug conditions, revealing major single-cell phenotypic populations and treatment-associated shifts in morphological state. Right: projection of averaged condition-level feature profiles, emphasizing drug-specific phenotypic shifts. (C) Left: hierarchical clustering dendrogram of drug conditions based on composite response scores integrating whole-spheroid volume, segmentation-based single-cell inhibition, and ATP-based CTG viability readouts. Right: PCA projection of composite response profiles with k-means clustering overlay, highlighting distinct phenotypic groups. Dactolisib and Temsirolimus tend to form separate clusters, while Pirfenidone, Epacadostat, and Lenalidomide tend to segregate into a distinct group. (D) Regional heatmaps showing drug effects on cancer cell nuclear area (left) and cytoplasmic area (right), segmented by spheroid regions (core, middle, periphery). Adjacent volcano plots display $\log_2$ fold-changes and statistical significance ($-\log_{10}$ p-values) for each treatment versus DMSO control per region. (E) Cell density plots for cancer and immune cells across drug conditions, normalized by spheroid volume. Error bars indicate mean ± SEM.

**Supplementary Text A. Segmentation pipeline - Implementation details**

All computational analyses were performed using custom Python scripts and Jupyter notebooks (python script: https://github.com/bioimage-profiling/Bayesian-segmentation3D). Image preprocessing included brightness adjustment (multiplicative scaling of the cytoplasmic channel), CLAHE-based contrast enhancement (with tunable clip limits and histogram bins), and contrast stretching (based on percentile clipping and intensity scaling). These preprocessing steps were applied independently to the cytoplasmic and nuclear channels before normalization and segmentation.

Nuclear segmentation was performed using a pretrained StarDist3D model, with the implementation available through the project repository (https://github.com/itarampoulous/StarDist3D), while whole-cell segmentation was carried out using the loyal-squid 3D U-Net model available through BioImage.IO/Zenodo (https://doi.org/10.5281/zenodo.7774505) with patch-wise inference. Custom tiling and halo configurations enabled processing of large 3D volumes. Incompatible tiling–halo combinations that violated image size or architectural constraints were automatically detected and skipped. For such cases, zero-filled fallback masks were returned to ensure uninterrupted optimization.

Final cell masks were generated using marker-based watershed segmentation, where StarDist-derived nuclei were used as seeds and cytoplasmic boundaries from the U-Net model served as constraints. These masks were evaluated using a composite objective function incorporating multiple segmentation quality metrics.

**Image formatting**

All images were processed in their native bit depth (typically 16-bit TIFFs) unless model requirements specified otherwise. Prior to segmentation with StarDist3D, images were normalized to the [0, 1] range using percentile-based scaling. CLAHE and contrast stretching were applied directly to the original intensity values without conversion to 8-bit. Output masks were saved in 16-bit format unless explicitly downsampled for visualization or export.

**Tiling and halo compatibility in 3D segmentation**

During patch-wise inference with 3D U-Net models, it is essential to ensure that the selected tile and halo sizes are compatible with both the image dimensions and the network architecture. Incompatible configurations may lead to segmentation artifacts or runtime errors. For each spatial axis (x, y, z), the following conditions must be met:

- Fit constraint:

$$\text{tile size} + 2 \times \text{halo size} \leq \text{image dimension}$$

- Divisibility constraint:

$$(\text{tile size} + 2 \times \text{halo size}) \mod 2^d = 0$$

where *d* is the number of downsampling layers (typically *d*=4).

These rules ensure compatibility with the receptive field and pooling structure of the model. All proposed tiling–halo configurations were validated against these conditions and excluded if invalid. A *utility script for verifying tile–halo compatibility is provided in the GitHub repository.*

**Supplementary Text B. Protocol for High-Resolution Single-Cell Profiling in 3D Spheroid Cultures Using MorphoNavigator-3D**

This protocol outlines the complete wet-lab and dry-lab workflow for single-cell resolution analysis of 3D spheroids, optimized for renal cancer (786-O) monocultures and T-cell co-cultures. It includes dye-based labelling, drug treatment, 3D imaging, ATP-based viability and flow cytometry assays, and automated single-cell segmentation using Bayesian optimization.

**Part 1: Cell culture and pre-labelling**

**1.1. Cell types and seeding**

- **Cancer cells**: 786-O clear cell renal cell carcinoma cells
- **Immune cells**: JE6.1 (Jurkat T-cells)
- **Plate**: Ultra-low attachment 384-well U-bottom plates (Corning, CLS3830)

| Culture Type | Seeding Density |
|---|---|
| **Monoculture (786-O)** | 500 cells/well |
| **Co-culture (786-O + JE6.1)** | 500 + 250 cells/well |

**1.2. Cytoplasmic pre-labelling**

Label cells **before seeding** using CellTracker dyes:

| Cell Type | Dye | Concentration | Incubation |
|---|---|---|---|
| **786-O** | CellTracker Orange | 2 μM (2X) | 45 min at 37°C |
| **JE6.1** | CellTracker Green | 0.5 μM (0.5X) | 45 min at 37°C |

- Wash cells twice with RPMI before seeding into wells (spin down for 5 min, 300 × g, then wash).
- Culture spheroids for 72 h in 25 µL RPMI-1640 supplemented with 10% FBS per well in 384-well ultra-low-attachment plates at 37 °C, 5% $CO_2$.

**Part 2: Nuclear staining and drug treatment**

**2.1. Live-cell nuclear staining (SPY650)**

- Add SPY650-DNA (1 µM final) 24 hours before imaging.
- Avoid 72 h incubation of SPY650 — causes spheroid degradation.
- Compatible with live imaging and downstream viability assays.

**2.2. Drug treatment**

- Add drugs directly to wells at time of cell seeding (day 0) or after 24 h of culture.
- Incubate for 72 h total drug exposure.
- Example drugs: Dactolisib, Temsirolimus, Epacadostat, Lenalidomide, Pirfenidone.
- Drugs diluted with 0.1% DMSO used as the vehicle control.
- Use (preferably) at least 3 concentrations (e.g., 1 nM, 100 nM, 1000 nM).

**Part 3: Endpoint assays**

**3.1. ATP-based cell Viability: CellTiter-Glo (CTG)**

- Add 15 µL CellTiter-Glo® 2.0 reagent (Promega, Cat. #G924) per well.
- Shake plates for 5 min, centrifuge at 300 × g for 3 min, and incubate at room temperature for 30 min to allow the luminescent signal to stabilize.
- Measure luminescence using a PHERAstar FS microplate reader (BMG Labtech).
- Normalize luminescence values to DMSO vehicle controls.

**3.2. Flow Cytometry (optional validation)**

- Add 50 µL of 0.25% Trypsin-EDTA per well.
- Incubate at 37°C with gentle pipetting (5–45 min). Pipette 10 times every 5-10 minutes, and keep incubating, until only separate single cells are observed.
- Quench with RPMI + FBS; transfer to 96-well plate.
- (optional) Stain with DRAQ7 (3 µM) for dead cell exclusion.
- Analyze on iQue Plus cytometer; gate by CellTracker dyes.

**Part 4: Imaging and data acquisition**

**4.1. 3D image acquisition**

- **Microscope**: Opera Phenix (20x water immersion, NA 1.0).
- **Excitation**: 488 nm (JE6.1), 561 nm (786-O), 640 nm (nuclei), and 405 nm when applicable.
- **Settings**: Spinning-disk confocal imaging with approximately 45 z-planes acquired at 2.5 µm intervals.
- Export 3D stacks in .TIFF format for downstream processing.

**Part 5: Image analysis pipeline**

**5.1. Preprocessing**

**Examples used in the optimization pipeline:**

- **Merge channels**: Cytoplasmic (orange/green) + nuclear (SPY650) → mean merge.
- **Brightness scaling**: Apply value ~10 to balance signal.
- **Contrast enhancement**: Use CLAHE or contrast stretching.

**5.2. Segmentation**

- **Nuclei**: StarDist3D (pre-trained model)
- **Cells**: loyal-squid 3D U-Net model
- **Refinement**: Watershed with nuclear masks as seeds

**Part 6: Bayesian optimization**

**6.1. Parameters optimized**

- Brightness scaling
- CLAHE clip limit and histogram settings
- Contrast-stretching parameters
- Segmentation foreground threshold

Tiling and halo settings were evaluated during preliminary development and subsequently fixed for the Bayesian optimization runs reported here.

**6.2. Objective function components**

- % cells in valid volume range (900–8200 $\mu m^3$)
- % pixel coverage inside masks
- Cell extent (extent-based shape penalty)
- Penalty for low cell counts
- Run using gp_minimize() from scikit-optimize
- Typical run: 30–80 iterations with 10 evaluations per cycle

**Part 7: Feature extraction and spatial analysis**

**7.1. Whole-spheroid features**

- Total volume (from cancer fluorescence)
- Average intensity
- Surface area

**7.2. Single-cell features**

- Nuclear/cell volume, intensity
- Cell coordinates for distance metrics

- Morphology (extent, solidity, etc.)

**7.3. Regional analysis**

- Assign each cell to **core**, **middle**, or **periphery** based on distance from spheroid centroid (3D Euclidean)

**7.4. Composite phenotyping**

- Compute and combine viability, cell morphology, spatial dispersion, immune engagement, etc.
- Normalize and visualize using radar (circular) plots

**Adaptation tips**

- Substitute other cell types with similar seeding and labelling logic.
- For microscopic imaging of live or fixed cells, SPY650 can be replaced by DRAQ5 or similar.
- For multi-channel phenotyping, adjust imaging setup accordingly.
- Flow cytometry can complement image-based assessment of cell proportions, although differences introduced by spheroid dissociation and sample processing should be considered.
